# Connectivity at fine scale: mapping structural connective fields by tractography of short association fibres *in vivo*

**DOI:** 10.1101/2024.04.30.591798

**Authors:** Fakhereh Movahedian Attar, Evgeniya Kirilina, Luke J. Edwards, Daniel Haenelt, Kerrin J. Pine, Robert Trampel, Denis Chaimow, Nikolaus Weiskopf

## Abstract

The extraordinary number of short association fibres (SAF) connecting neighbouring cortical areas is a prominent feature of the large gyrified human brain. The contribution of SAF to the human connectome is largely unknown because of methodological challenges in mapping them. We present a method to characterise cortico–cortical connectivity mediated by SAF in topologically organised cortical areas. We introduce the ‘structural connective fields’ (sCF) metric which specifically quantifies neuronal signal propagation and integration mediated by SAF. This new metric complements functional connective field metrics integrating across contributions from short- and long-range white matter and intracortical fibres. Applying the method in the human early visual processing stream, we show that SAF preserve cortical functional topology. Retinotopic maps of V2 and V3 could be predicted from retinotopy in V1 and SAF connectivity. The sCF sizes increased along the cortical hierarchy and were smaller than their functional counterparts, in line with the latter being additionally broadened by long-range and intracortical connections. *In vivo* sCF mapping provides insights into short-range cortico– cortical connectivity in humans comparable to tract tracing studies in animal research and is an essential step towards creating a complete human connectome.

**Highlights:**

- Non-invasive mapping of Short Association Fibre (SAF) connectivity via diffusion-weighted MRI-based probabilistic tractography accurately predicted cortical functional neuroanatomy.
- The novel structural Connective Fields (sCF) concept provides a quantitative measure of cortico-cortical integration facilitated by SAF, complementing the existing functional Connective Field (CF) concept.
- Sub-millimeter resolution diffusion-weighted MRI enables tractography and connective field modeling of SAF, unlocking applications previously restricted to invasive tract tracing in animal studies.

## Introduction

Understanding how the complex connectivity of cortical neurons determines their responses to sensory stimuli and facilitates cortical computations is a fundamental challenge in neuroscience. In the primate visual cortex, fast and efficient feedforward and feedback connections from the primary visual cortex (V1) onward enable rapid processing of complex visual information. Neurons in V1 encode local image features, while higher-order visual areas like V2 and V3 respond to more distributed and complex image features in a hierarchical, recurrent process [1–7] (Fig. 1a,b). Spatial integration of visual information is enabled by a complex network of intracortical fibres within each area as well as by converging and diverging white matter connections between the hierarchy levels [8] (Fig. 1c). Each type of connection plays a specific role in the generation of cortical responses but their relative contributions are not precisely known, especially in humans.

**Fig. 1:**
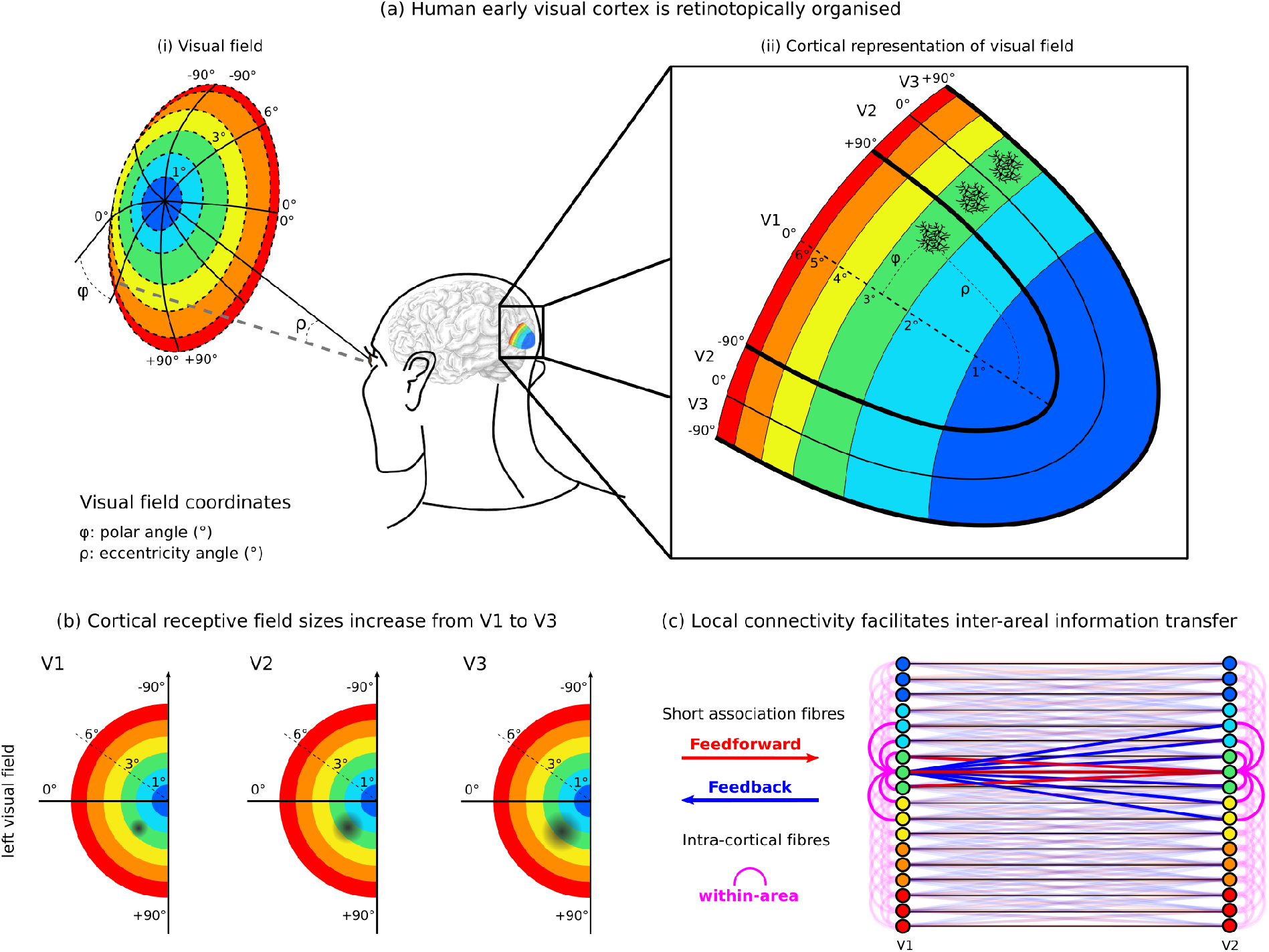
Retinotopic organisation of visual cortical areas and information transfer from the visual field to the cortical surfaces of V1, V2 and V3. (a) Each point in (i) the visual field and (ii) its cortical representation is encoded using two coordinates: polar angle (*φ*) and eccentricity (*ϱ*). (b) The receptive field sizes of cortical neurons increase from V1 to V3, represented by the increasing size of the black patch. (c) Information transfer and integration between and within different levels in the visual hierarchy is enabled by converging feedforward and diverging feedback white matter connections and intracortical fibres. Together, all these connections determine the cortico–cortical connective fields of cortical neurons and facilitate integration of visual information.

A striking feature of the large and gyrified human brain is the large number of short association fibres (SAF) in the superficial white matter connecting directly adjacent cortical areas. SAF make up almost 90 % of all white matter connections in the human brain [9]. They mediate local cortico–cortical connectivity by linking different cortical hierarchy levels [2, 10] over distances of up to approximately 30 mm [9].

In primate studies, the distinct role of SAF in cortical integration within the early visual processing stream has been demonstrated by invasive electrophysiology, tracer injections and functional optical imaging [6, 11–15].

In the human brain, SAF are largely unexplored mainly because of technical challenges in mapping them. For example, it is not known to what extent the integration of visual information in a higher cortical area is realised by converging SAF from a lower area and to what extent by intracortical fibres. Direct transfer of findings from non-human primates is not readily possible because the substantial expansion of the cortex and the unique sulcal anatomy of the human brain [16] result in different geometry and sizes of SAF. Functional MRI (fMRI) can be used to map cortical population receptive fields [3, 17] and connective fields [5, 18, 19] as functional measures of cortical integration along the visual processing hierarchy *in vivo*. However, functional measures do not provide information about the white matter underpinnings of cortical integration. This leaves diffusion-weighted imaging (DWI) tractography [20] as the only tool to non-invasively study the SAF connections and their detailed organisation in humans, where invasive tracer studies are not applicable.

Here, we present a new approach that combines sub-millimetre spatial resolution DWI for tractography of SAF with fMRI for retinotopic mapping to specifically map and characterise cortico–cortical connectivity mediated by SAF in the human early visual processing stream. The method could also be applied in other topologically organised cortical areas including motor, somatosensory and auditory cortices. It builds on a recently developed and validated multi-modal MRI approach that addressed several key technical challenges of mapping the SAF *in vivo* by sub-millimetre spatial resolution DWI tractography [21].

In order to quantify the specific contribution of SAF to information transfer and cortical integration between different levels of cortical hierarchy we introduce a new metric – structural connective fields (sCF). This metric complements the established functional connective field metric obtained using fMRI, which encompasses and mixes multiple contributions from intracortical as well as short- and long-range white matter connections.

We found highly coherent spatial arrangements of SAF preserving cortical functional topology up to V3. The retinotopy in higher cortical areas (V2 and V3) could be predicted with high accuracy and precision based solely on their SAF connections to V1 and V1 retinotopy, particsularly along the cortical direction aligned with eccentricity encoding. The sCF metric allowed us to estimate cortico–cortical sampling sizes mediated by SAF, which increased along the visual hierarchy in line with known functional connective field sizes. Along the cortical direction aligned with polar angle encoding, sCF predicted the functional connective field characteristics less well, due to the gyral bias [22, 23] of SAF detectability in tractography.

### Structural connective field mapping

We introduce the structural connective fields (sCF) framework in analogy to the established functionally-defined connective fields framework. In particular, we target the structural measures of cortico–cortical integration in the human early visual processing stream. In this section, we first summarise the functional connective fields metrics between different cortical hierarchy levels (V1–V2 and V1–V3) obtained in previous fMRI studies in the human brain. We then introduce a novel method to map the sCF based on probabilistic tractography of SAF combined with fMRI retinotopic mapping *in vivo*. The detailed theory of the structural connective field framework and the description of retinotopic processing of visual information in the early visual cortex are given in Appendix A.

### Functional cortical connective fields estimated from fMRI

The early visual cortex is retinotopically organised [18, 24–28]. This means that adjacent points in the visual field are encoded by neighbouring groups of neurons on the cortical surfaces of V1, V2 and V3. This organisational principle also imposes the two-dimensional visual field coordinate system defined by eccentricity (*ϱ*) and polar angle (*φ*) onto the surface of each early visual cortical area (Fig. 1a). The visual field coordinates (*ϱ* and *φ*) can be mapped onto each location in V1, V2 and V3 using fMRI retinotopic mapping [26, 27].

The connective fields framework in the early visual processing stream describes information transfer and integration between neuronal populations of V1, V2 and V3 in retinotopically corresponding locations. For a pair of cortical areas, the connective field defines the cortical location and extent in one area connected to a particular location in another area. Connective fields between early visual cortices have been estimated with fMRI using the connective fields framework based on the analysis of aggregate functional activity [5]. The estimated connective fields reflect overall information integration comprising feedforward and feedback inter-area SAF as well as intracortical intra-area connections. For each location in the higher area *V*_*n*_ the connective field is typically determined as a two-dimensional Gaussian filter on the cortical surface of an earlier area *V*_*m*_ (*m < n*) that can optimally describe the temporal evolution of the cortical signal in *V*_*n*_. The peak of the Gaussian determines the cortical location in *V*_*m*_ with highest connectivity to *V*_*n*_ and its width gives a measure of the connective field size. Connective field sizes obtained using this method are approximately 3 mm for V1–V2 connections and increase along the visual processing stream, reaching 5–6 mm for V1–V3 connections [5].

### Structural connective field sizes estimated with SAF DWI tractography

We introduce the structural connective fields (sCF) framework which follows the functional connective fields concept, but quantifies the specific contributions of SAF to cortico–cortical information transfer and integration. We define the sCF as the normalised distribution of SAF termination densities on the cortical surface of a lower area *V*_*m*_ connected to a particular neighborhood *x*_*n*_ in a higher area *V*_*n*_ (*m < n*). The centre of mass of the sCF determines the cortical location in the lower area with highest SAF connectivity to *x*_*n*_. The width of the distribution defines the sCF size and gives a measure of the cortico–cortical integration realised specifically by SAF.

Assuming that the spatial distribution of SAF streamline terminations in the cortex mapped by probabilistic DWI tractography accurately reflects the underlying spatial distribution of SAF connections, we estimated the sCF by combining the SAF streamlines with visual field coordinates of their cortical terminations – eccentricity (*ϱ*) and polar angle (*φ*) – obtained by fMRI retinotopic mapping (Fig. 2a,b).

**Fig. 2:**
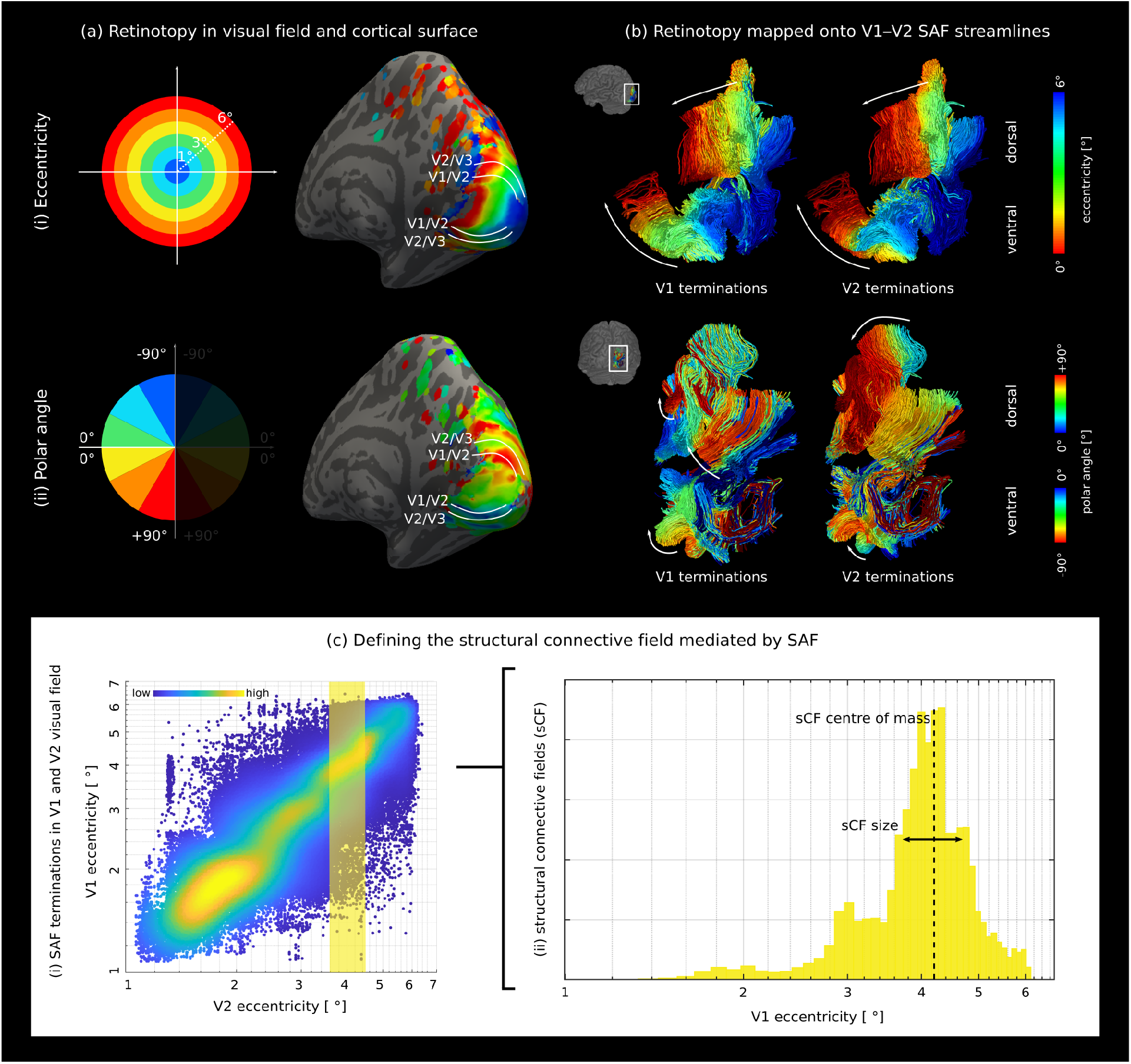
Structural connective fields (sCF) mapped by combining retinotopic maps from fMRI and SAF from DWI probabilistic tractography on a fine scale. V1–V2 SAF are shown for a representative hemisphere. (a) Smoothed eccentricity (*ϱ*) and polar angle (*φ*) maps were (b) sampled onto SAF streamlines connecting V1–V2, assigning two pairs of visual field coordinates (*ϱ*_*V*1_, *φ*_*V*1_) and (*ϱ*_*V*2_, *φ*_*V*2_) to each streamline. A topological spatial arrangement of SAF was observed for both coordinates *ϱ* and *φ* (white arrows). Illustrated here for V1–V2 eccentricity, (c) the sCF is defined as the histogram distribution of visual field coordinates of SAF cortical terminations in V1 projected from a defined region (highlighted in (i)) in the visual field representation of V2. Here, the histogram presents the group–level analysis. The sCF centre of mass represents the most strongly connected locations between the cortical areas and describes the cortico–cortical information transfer, while the sCF size is determined as the width of the distribution and describes the contribution of SAF to cortico–cortical integration. See Appendix A for details of sCF mapping theory.

To characterise the sCF, the visual field coordinates corresponding to the cortical terminations of SAF streamlines were analysed as exemplified for V1–V2 connections in Fig. 2c. Each point on the scatter plot depicts an individual streamline connecting V1 (or another lower area *V*_*m*_) and V2 (or another higher area *V*_*n*_) shown on the vertical and horizontal axes, respectively. For streamlines terminating in a small neighbourhood *x*_*n*_ in the higher area *V*_*n*_ (e.g. the highlighted area in Fig. 2c(i)), the histogram of the visual field coordinates mapped to their terminations in a lower area *V*_*m*_ (Fig. 2c(ii)) illustrates the distribution representing the sCF. The peak of the distribution, or its centre of mass, describes the average location in *V*_*m*_ to which *x*_*n*_ in *V*_*n*_ is connected and its width is used as an estimate of sCF size.

## Results

We applied a novel metric – structural connective fields (sCF) – to specifically quantify cortico–cortical connectivity mediated by SAF in the human early visual processing stream. For this purpose, we conducted a fine-grained analysis of connectivity among three pairs of early visual cortical areas (V1–V2, V2–V3, and V1–V3). We utilised sub-millimetre spatial resolution probabilistic DWI tractography facilitated by the high performance gradient system of the 3 T Connectom scanner to identify the sCF connecting different levels of the cortical hierarchy. Our data consisted of 18 hemispheres from 11 healthy participants (using a subset of data from our previously published study [21] where V3 was also included in the DWI field-of-view). The data were combined with fMRI retinotopic maps from the same individuals to determine the visual field coordinates corresponding to SAF cortical terminations.

First, to showcase the efficacy of the new metric in describing cortico–cortical information transfer, we demonstrated that visual field coordinates in a higher cortical area can be predicted based on its white matter connectivity to a lower area using the computed centre of mass of the sCF.

Second, to illustrate the utility of the new metric to characterise cortico–cortical integration, we computed the sCF sizes for each pair of early visual cortical areas. We demonstrated the expected increase in cortico–cortical sampling sizes at higher levels of the cortical hierarchy, in line with functional connective field mapping.

Finally, we related structural and functional measures of cortico–cortical connectivity by comparing our estimates of sCF size to prior fMRI-estimated connective field sizes. This analysis allowed us to quantify the contribution of sCF connectivity to cortico–cortical integration and evaluate the effectiveness of our sCF mapping approach.

### SAF tractography predicts functional retinotopy *in vivo*

A dense network of SAF interconnecting V1, V2, and V3 were identified in all the studied hemispheres (see example of V1–V2 SAF in Fig. 2b) by sub-millimetre spatial resolution DWI probabilistic tractography optimised for SAF detection [29]. Highly coherent retinotopic spatial arrangements of SAF in white matter (Fig. 2b) were revealed by projecting the visual field coordinates of their cortical terminations along the SAF pathways. Notably, SAF terminating in cortical locations encoding neighbouring areas of the visual field displayed closely aligned trajectories within the white matter (Fig. 2b).

Retinotopic organisation of SAF connectivity was corroborated by quantitative analysis combining streamlines from tractography and cortical visual field coordinates from fMRI retino-topic mapping (Fig. 3). Group-level findings are presented in a scatter plot (Fig. 3) where each data point depicts an individual streamline. The horizontal and vertical axes correspond to the visual field coordinates of SAF cortical terminations in the higher (*V*_*n*_) and the lower (*V*_*m*_) visual cortical areas, respectively. Retinotopic organisation implies that strongest SAF connectivity will be between cortical locations encoding the same visual field locations at different levels of the cortical hierarchy, implying that the strongest connections should lie along the line of identity (not shown) in the scatter plot.

**Fig. 3:**
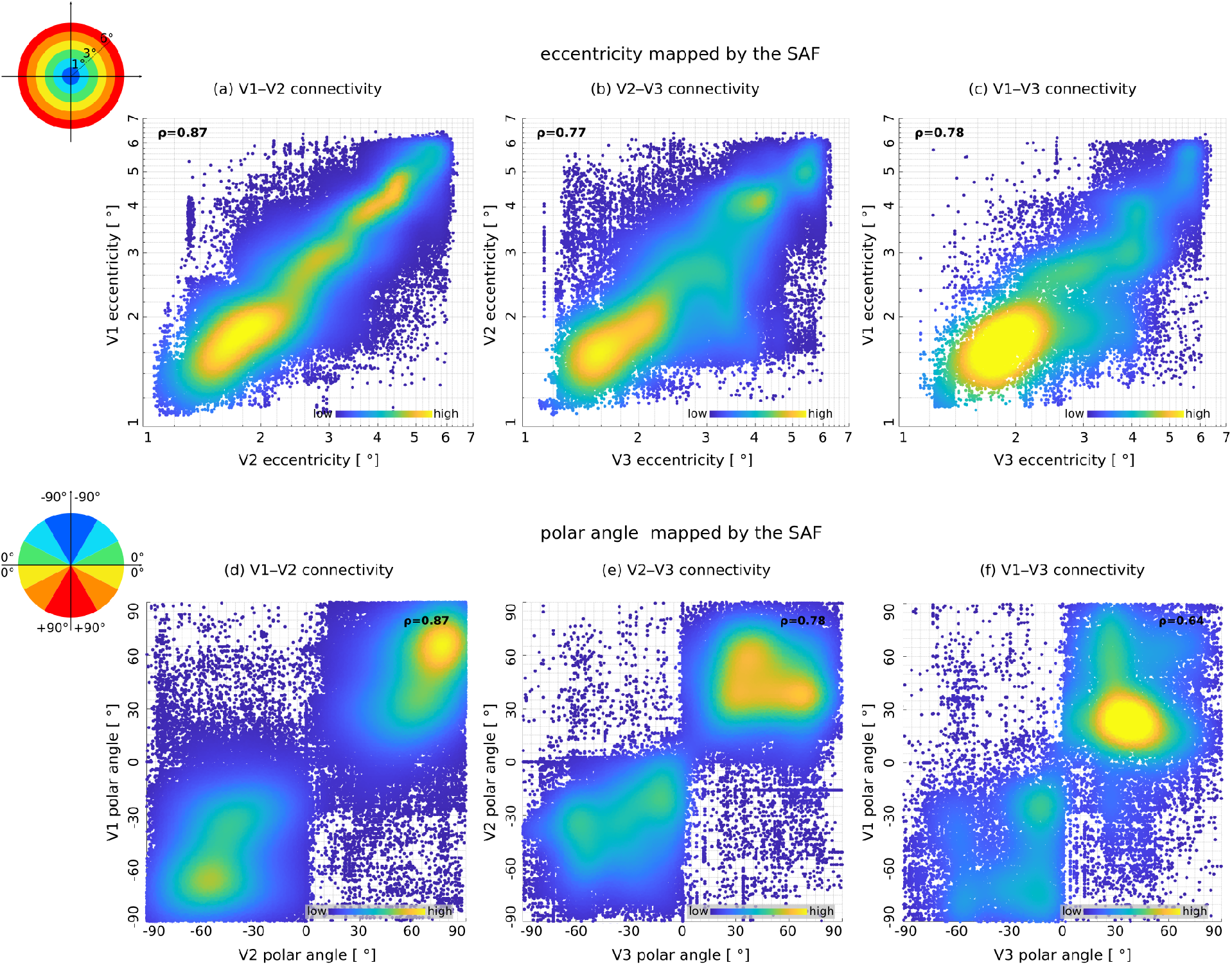
SAF connectivity mapped by probabilistic tractography preserved cortical retinotopy. Density plots illustrate eccentricity (*ϱ*) (a-c) and polar angle (*φ*) (d-f) mapped to SAF streamline terminations connecting V1–V2, V2–V3 and V1–V3 over all hemispheres. The eccentricity plots were pooled over all polar angles, and the polar angle plots were pooled over all eccentricities. Our fMRI experiment covered approximately 6^*0*^ in the visual field eccentricity direction. The polar angle was parameterised to the range [−90^*0*^, 90^*0*^] in both hemifields based on its deviation from the horizontal meridian, where positive values represented the upper visual field, enabling averaging across the left and right hemispheres. Each point on the scatter plot corresponds to one streamline, with its visual field coordinate in *V*_*m*_ on the y-axis and *V*_*n*_ (*m< n*) on the x-axis. The colour scale shows the point density.

In line with these expectations, the eccentricity coordinates of SAF terminations for all pairs of cortices V1–V2, V2–V3 and V1–V3 demonstrated a high Pearson’s correlation coefficient (*ρ*) computed over all the streamlines (*ρ* = 0.87, 0.77 and 0.78 for V1–V2, V2–V3 and V1–V3 connections, respectively). This was also reflected in the higher density of data points around the *ϱ*_*n*_ = *ϱ*_*m*_ line of identity in Fig. 3a-c, especially for the V1–V2 connections. Higher density was observed for low eccentricity values, which corresponded to the representation of the visual field centre.

Less coherent fibre arrangements were found in the polar angle direction (*φ*) (Fig. 2b(ii), Fig. 3d-f). If both the upper (+) and lower (−) visual fields were considered together (Fig. 3d-f), a high Pearson’s correlation coefficient for V1–V2 and V2–V3 connections (*ρ* = 0.87 and 0.78, respectively) and only a moderate one for V1–V3 connections (*ρ* = 0.64) was observed. However, within the upper and lower visual fields separately, where most of the streamlines were detected, the computed correlations dropped to much smaller values (*ρ* (−/+) = 0.30/0.36, 0.00/0.25 and 0.00/0.14 for V1–V2, V2–V3 and V1–V3, respectively) pointing to relatively low apparent retinotopic order in the polar angle direction within the cortical representation of each sub-field of the visual field.

The retinotopic organisation of SAF, especially along the cortical direction aligned with eccentricity encoding (Fig. 3a-c), opens up the opportunity to predict visual field coordinates in a higher cortical area based solely on its SAF connectivity to a retinotopically mapped lower cortical area using its sCF centre of mass. This is qualitatively demonstrated for SAF connectivity between the three pairs of cortices at the group level on a template brain surface (Fig. 4) and at the single-subject level in each individual’s native space (Supplementary Figs. 8, 9 and 10). At the group level, the predicted visual field coordinates at each location in a higher cortical area were determined as its sCF centre of mass by averaging the visual field coordinates of all SAF projecting to this location from a lower area.

**Fig. 4:**
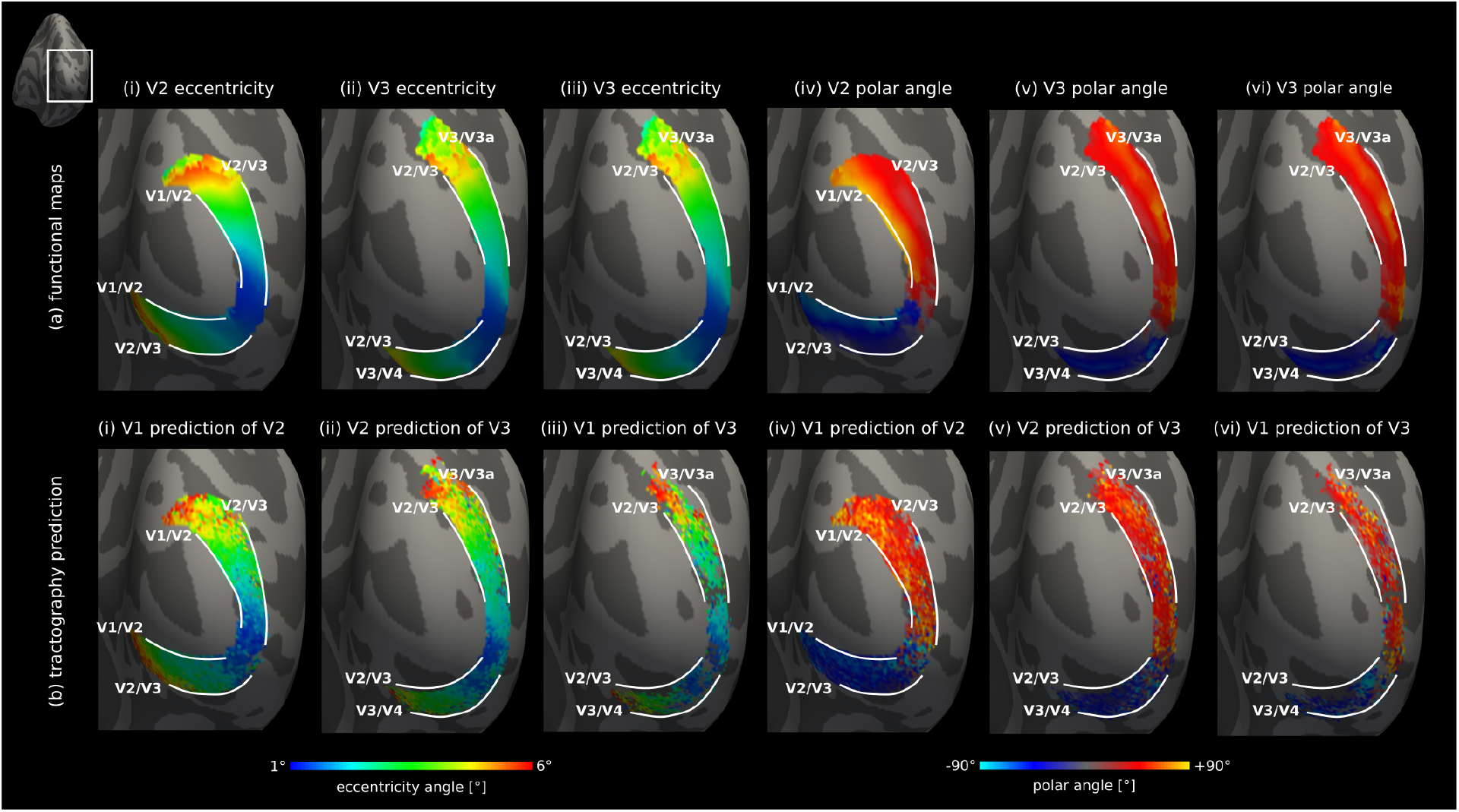
Retinotopic order in a higher visual area can be predicted based on its SAF connectivity to a lower visual area. Group-averaged maps over 11 right hemispheres projected onto the inflated surface of the fsaverage (FreeSurfer [30]) template. (a) Visual field coordinates – eccentricity and polar angle – in V2 and V3 determined by fMRI and (b) their corresponding tractography-generated predictions based on SAF connecting (i,iv) V1–V2 and V1 retinotopy, (ii,v) V2–V3 and V2 retinotopy, and (iii,vi) V1–V3 and V1 retinotopy. FreeSurfer atlas labels were used to determine the V1, V2 and V3 areal borders on the template surface.

The resulting connectivity-based prediction of retinotopic maps in V2 and V3 (Fig. 4b, Supplementary Figs. 8b, 9b and 10b) closely resembled retinotopic maps obtained directly and independently using fMRI (Fig. 4a, Supplementary Figs. 8a, 9a and 10a). The most accurate results were obtained for visual field coordinates in V2 based on V1–V2 SAF connectivity, which also have the shortest geodesic distances (retinotopic SAF lengths of 12.7 *±* 5.6 mm (mean*±*standard deviation)). Shorter fibres are expected to benefit from enhanced tractography performance relative to longer fibres. The least accurate results were obtained for V3 based on V1–V3 SAF connectivity with retinotopic SAF lengths 25.0*±*7.3 mm (mean*±*standard deviation), possibly reflecting the larger errors in tractography for these fibres. Prediction was more accurate for the eccentricity compared to the polar angle encoding direction, likely because of tractography gyral biases affecting the results in the latter (see section on gyral bias effects below).

The prediction of cortical retinotopy based on SAF connectivity was also possible at the individual participant level (Supplementary Figs. 8, 9 and 10). Similar to the group-analysis (Fig. 4b), the most accurate predictions were obtained for retinotopy in V2 based on V1–V2 connectivity. Also, more accurate results were obtained for V2 and V3 based on V1–V2 and V2–V3 connectivity in the given order in line with the group results, especially in the eccentricity encoding direction. The overall poorest performance was observed for V3 based on V1–V3 connectivity in the polar angle encoding direction, which may be explained by both distance and gyral bias effects. The patterns of SAF streamline terminations on the cortical surface in individual hemispheres revealed uneven coverage, with particularly sparse coverage for higher cortical area V3. Inter-individual differences in cortical gyrification patterns resulting in different gyral bias patterns likely differentially affected the tractography performance per hemisphere, resulting in varying patterns of SAF cortical terminations (Figs. 8b, 9b and 10b). These effects were partly averaged out at the group level.

Together, these results demonstrate the utility of high-resolution SAF tractography to map the sCF centre of mass, enabling neuronal signal propagation to be tracked between the cortical hierarchical levels at a fine spatial scale.

### sCF size increases along the visual hierarchy

The sCF size was estimated to quantify cortico–cortical integration between different levels of the visual cortical hierarchy mediated specifically by SAF. We estimated the sCF size for V2 and V3 for the SAF connecting the three pairs of cortical areas V1–V2, V2–V3 and V1–V3. It was calculated as the standard deviation of the distribution of differences in the visual field coordinates of SAF cortical terminations in the lower area *V*_*m*_ and the higher area *V*_*n*_, binned in *V*_*n*_ over bins with width of 1^*°*^ (Supplementary Fig. 11). The estimates were obtained over all the streamlines in *V*_*m*_ connected to these small neighborhoods in *V*_*n*_ across all the studied hemispheres (see Appendix A). We determined the sCF sizes along the cortical directions aligning with eccentricity and polar angle directions separately, each expressed first in visual field units (Supplementary Fig. 12) and then translated to distances on the cortical surface measured in millimetres (Fig. 5).

**Fig. 5:**
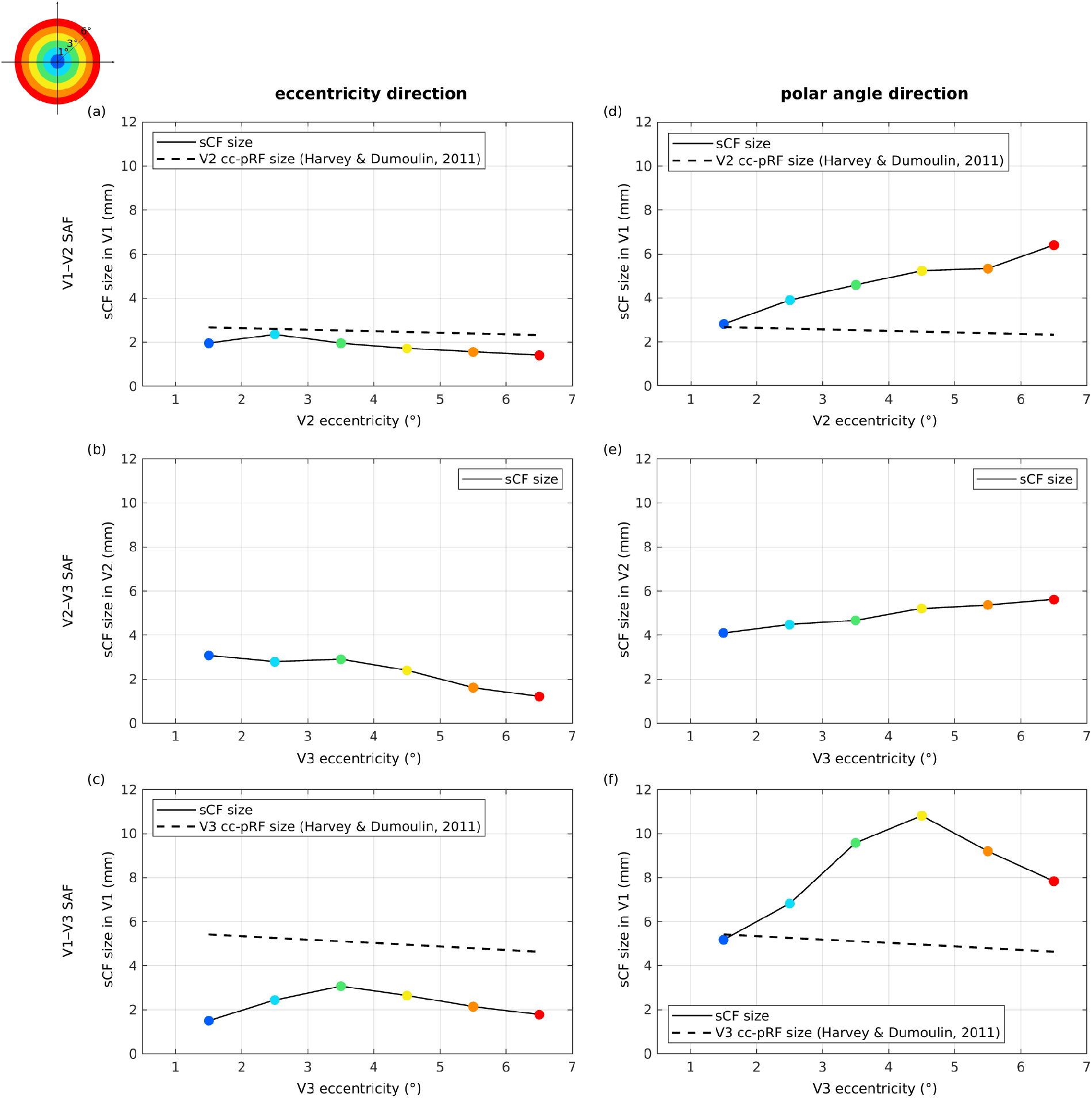
Estimated sCF sizes (mm) for cortical directions aligned with eccentricity (a-c) and (d-f) polar angle encoding. Group-level estimates are shown for V1–V2, V2–V3 and V1–V3 connections as black solid lines connecting the coloured points. The sCF size was estimated as the standard deviation of the distribution of the differences in the visual field coordinates of SAF terminations sampled on the cortical surface in *V*_*m*_ and *V*_*n*_. For both eccentricity and polar angle, the sCF size was calculated for *V*_*m*_–*V*_*n*_ SAF terminating in 1^*°*^ bins of visual field eccentricity in *V*_*n*_ (*m < n*) and was translated into cortical surface units by the cortical magnification factors in *V*_*m*_ (see Appendix A). Each coloured point corresponds to the sCF size of one bin. For comparison, literature values of fMRI-estimated cortico-cortical sampling sizes (cc-pRF) from Ref. [3] are plotted as dashed lines for sampling of *V*_*m*_ in *V*_*n*_ (see Appendix A for definition of cc-pRF size). That work did not report cc-pRF sizes for V2-V3 connections, hence not shown.

The estimated sCF sizes (measured as distances (mm) on the cortical surface) increased up to V3, consistent with fMRI-estimated connective field sizes [3, 5], reflecting the higher degree of visual integration at higher levels of the cortical hierarchy. We found larger sCF sizes for V1–V3 and V2–V3 connections compared to V1–V2 connections along both cortical directions (Fig. 5).

For the eccentricity encoding direction, our average estimates of the sCF size (mean*±*standard deviation) on the cortical surface were 1.8 *±* 0.3 mm for V1–V2, 2.3 *±* 0.8 mm for V2–V3 and 2.3 *±* 0.6 mm for V1–V3 SAF (Fig. 5a-c). For this direction, sCF sizes were nearly constant for V1–V2 SAF (Fig. 5a) but slightly decreased towards the cortical locations representing the periphery of the visual field, also observed to a greater extent for V2–V3 SAF (Fig. 5b). This finding was in line with the known decrease in the functional cortico-cortical sampling sizes in the periphery of the visual field [3] as shown by a dashed line on the graph (Fig. 5a,c). Expressed in visual field angle (^*°*^) units, the sCF sizes measured along the eccentricity encoding direction slightly increased towards the visual field periphery for V1–V2 SAF and were about 0.5^*°*^, in line with the fMRI-estimated connective field sizes and population receptive field measures, which are known to also continuously increase towards the visual field periphery (Supplementary Fig. 12a-c). For V2–V3 and V1–V3 SAF, the sCF sizes were approximately constant towards the periphery and were about 0.7^*°*^. Estimates close to the visual field centre and on the outer periphery of the mapped visual field may be affected by artifacts from border effects in fMRI retinotopic mapping experiments.

When compared to fMRI-estimated connective field sizes reported in the literature [3, 5], sCF sizes measured along the eccentricity encoding direction were comparable (for V1–V2 connections) or smaller (for V1–V3 connections) than the functional measures of 2.5 *±* 0.1 mm for V1 sampling in V2 (or V2 cc-pRF size) and 5.0 *±* 0.3 mm for V1 sampling in V3 (or V3 cc-pRF size)[3, 5]. (Fig. 5a,c). This observation is neuroanatomically plausible, as functional connective field sizes are additionally influenced by intracortical, feedback SAF and long-range connections and, therefore, expected to be larger. The neuroanatomical plausibility of our sCF size estimates in the cortical direction aligned with eccentricity encoding suggests only a small contribution of measurement errors (see Appendix A). Together, these findings corroborate the utility of the proposed sCF framework to quantify cortico–cortical integration and neuronal signal propagation by tractography of SAF.

However, the estimated sCF sizes were much larger in the cortical direction aligned with polar angle (Fig. 5d-f) than along eccentricity encoding. Our average estimates (mean*±*standard deviation) were 4.7 *±* 1.3 mm for V1–V2 and 4.9 *±* 0.6 mm for V2–V3, and increased to around 8.2 *±* 2.1 mm for V1–V3 SAF (Fig. 5d-f). The sCF sizes estimated in the polar angle direction were also substantially larger than literature values of fMRI-estimated connective fields (dashed lines in Fig. 5d,f), suggesting they were overestimated by our method and point to a substantial contribution of tractography errors to our estimates (see below). Further, unlike the fMRI estimates, the sCF sizes increased towards the periphery of the visual field corresponding to a sCF size of approximately 0.5^*°*^ near the visual field centre to more than 3^*°*^ on its periphery (Supplementary Fig. 12d-f).

### Gyral bias of tractography affects sCF estimates

We analysed the potential impact of gyral bias on our SAF tractography in order to understand the poorer performance of our method to estimate the sCF sizes for polar angle encoding. In the human visual cortex, eccentricity encoding corresponds on the cortical surface roughly to lines running along the gyral crowns and sulcal fundi. This means that cortical regions representing different eccentricities, but same polar angles, have a similar position with respect to gyri and sulci and are, therefore, affected in a similar way by a gyral bias.

In contrast, polar angle encoding is approximately perpendicular on the cortical surface to the line representing eccentricity encoding. In this direction, regions representing different polar angle locations are located at different sulcal depths on the cortical surface and may be differentially affected by a gyral bias. Fig. 6a,b shows the density of V1–V2 SAF terminations in the cortex computed at the group level in a template space. Higher density was found around the gyri representing the vertical meridians of the visual field. This was also demonstrated in Fig. 3d where higher SAF densities were observed around the representations of the vertical meridians in the dorsal (near +90^*°*^) and ventral (near -90^*°*^) parts of the visual cortex. In contrast, much fewer SAF terminated in the calcarine sulcus, demonstrating the presence of a gyral bias that forced the SAF to preferentially terminate near the gyral crowns because of tractography errors. Thus, estimates of sCF size in the polar angle direction were also affected. This was reflected in sCF sizes averaging to about 30^*°*^ for V1–V2 and V2–V3, and 60^*°*^ for V1–V3 SAF, accounting for approximately 1/3 and 2/3 of the size of the cortical surface in the polar angle direction per visual field quadrant, respectively.

**Fig. 6:**
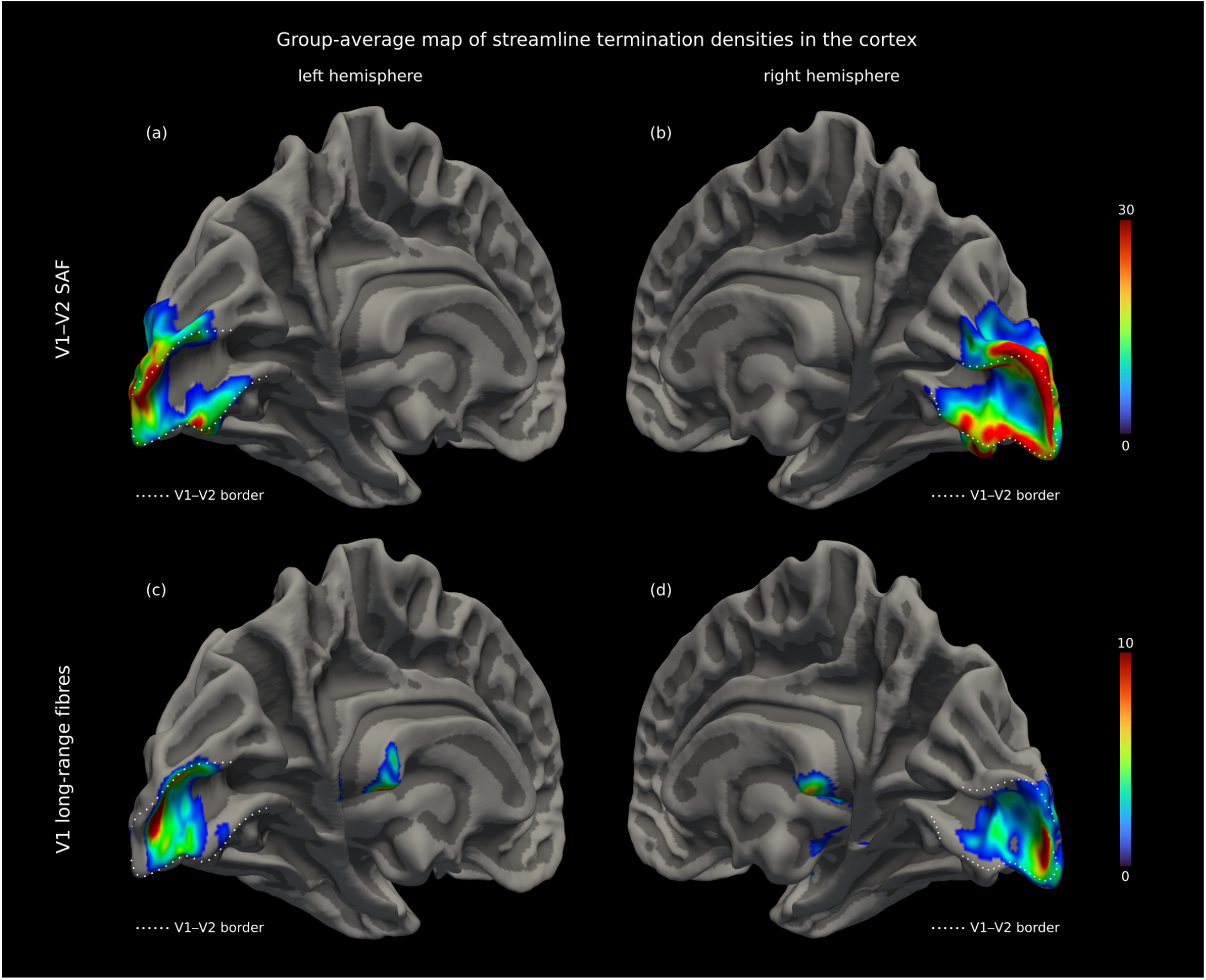
Gyral bias of tractography affects detectability of SAF near sulcal fundi. The spatial distribution of SAF termination densities in the cortex is shown for V1–V2 connections in the fsaverage (FreeSurfer [30]) template space for the (a) left and (b) right hemispheres. Higher density is observed on and near the gyral crowns compared to the walls and troughs of sulci, pointing to the presence of a gyral bias in the anatomical direction of polar angle encoding. (c,d) Results are also shown for the long-range fibre pathways terminating in V1, including the optic radiation, showing higher densities mainly near the sulci. The cortical termination patterns of SAF and long-range pathways are almost reciprocal (see Supplementary Fig. 13 for more detail).

The optic radiation, the major white matter bundle projecting to V1, showed cortical termination patterns almost reciprocal to SAF, showing higher density in the calcarine sulcus and lower density in the gyrus (Fig. 6c,d). The dominant fibres of the optic radiation may exacerbate the gyral bias of SAF detectability by deflecting tractography pathways and/or preventing SAF detection (see Supplementary Fig. 13 for more detail).

## Discussion

SAF are a major component of the human white matter connectome and understanding their roles in cortical processing is essential to understanding human brain function. This work developed a new method to non-invasively characterise information transfer and cortical integration specifically mediated by SAF in topologically organised cortical areas. By establishing a direct link between SAF and the functional neuroanatomy of overlying cortices, our method offers a new approach to explore the role of structural connectivity in cortical processing *in vivo*.

Our study demonstrates the capacity of the proposed method to finely map SAF connectivity, enabled by ultra-high spatial resolution DWI acquired using the high performance gradient system of the Connectom MRI scanner [31] and a dedicated data analysis approach. DWI methodology remains a rapidly developing field and new methods for ultra-high spatial resolution DWI may enable future translation of our method to widely available clinical scanners [32, 33].

We introduced a novel structural connective fields (sCF) metric to characterise the short-range white matter underpinnings of information transfer and cortical integration, complementary to the established functional connective field measures [3, 5, 19]. For each location on the cortical surface of a given area, the sCF provides the corresponding location on the cortical surface of another cortical area most strongly connected to it as well as the dispersion of their connecting fibres. In this way, the sCF describes the cortico–cortical sampling between neighbouring cortical areas. In cortical areas with topologic organisation, sCF can be directly related to information transfer and functional cortico–cortical integration of sensory stimuli. The sCF metric was designed in analogy to the connective field metric developed in the field of fMRI. While functional connective fields represent an effective measure of aggregate cortico–cortical connectivity irrespective of its structural underpinnings, sCF reflects specifically the cortico– cortical integration relayed by the SAF. It therefore approximates the connectivity measures derived from invasive tracer studies in animals.

We applied the proposed framework for sCF mapping to characterise SAF connectivity in the well-studied human early visual processing stream up to V3, but it can be readily applied to other brain areas. Below, we discuss the two main findings of our study and related important implications for its application across the human brain.

First, we mapped the SAF and demonstrated their highly coherent topological arrangements in the early visual processing stream. We demonstrated that retinotopic organisation in higher visual cortical areas V2 and V3 could be accurately predicted based on their direct SAF connections to V1, especially for the cortical direction aligned with eccentricity encoding (Fig.4). Therefore, tractography of SAF could in principle be used to explore topological organisation in hierarchically connected cortical areas across the brain based solely on their SAF connections and without the need for additional functional measures or electrophysiological recordings. This becomes especially important in applications where functional imaging is challenging, e.g. in blind individuals. Moreover, this finding provides a valuable direction for studying perceptual deficits caused by juxtacortical lesions, such as in multiple sclerosis [34] and stroke.

Second, cortico–cortical integration by SAF was characterised in the human brain by sCF sizes for the first time. Our estimates of the sCF size were in line with neuroanatomical knowledge, especially in the cortical direction aligned with eccentricity encoding. They increased along the visual hierarchy from V1–V2 to V1–V3, reflecting the increasing degree of visual integration in higher visual cortical areas as previously reported also by functional connective fields [3, 5]. The average (mean*±*standard deviation) sCF sizes were 1.8 *±* 0.3 mm for V1–V2 and 2.3 *±* 0.6 mm for V1–V3 in the eccentricity direction and the corresponding fMRI-estimated connective field sizes were 2.5 *±* 0.1 mm for V1 sampling in V2 and 5.0 *±* 0.3 mm for V1 sampling in V3 [3, 5]. Despite being comparable, our sCF sizes in the eccentricity direction were smaller than functional connective field sizes. This observation may point to the high coherence of SAF axons in superficial white matter, resulting in smaller sCF sizes. In the cortex, the additional contributions of disperse telodendritic trees as well as direct and indirect intracortical fibres may contribute to the larger estimated functional connective fields. Additionally, the functional connective fields estimates are blurred by hemodynamic spread [35, 36]. These additional contributions were more prominent for V1–V3 connections at higher hierarchy levels with average (mean*±*standard deviation) 5.0 *±* 0.3 mm connective fields [3] compared to 2.3 *±* 0.6 mm sCF sizes.

These results imply that although on a coarser spatial scale compared to tracer studies, the precision of our SAF probabilistic tractography enables similar questions to be addressed about the biological dispersion of fibres previously only possible with invasive tracer injections in animal studies. Therefore, it enables detailed studies of cortical processing also in humans non-invasively *in vivo*.

The successful application of our method in the early visual processing stream demonstrates its applicability to map brain topology on a fine-grained spatial scale. Previous studies in human [37–40] and macaque [41] also demonstrated the feasibility of DWI tractography to map white matter topography but only on a coarse spatial scale. Our method can be used to independently explore cortical topology, for instance, in the primary auditory cortex where tonotopic organisation is demonstrated along two orthogonal axes [42–45] or in the primary somatosensory cortex where somatotopic organisation is demonstrated along the central sulcus [46, 47]. It could also be applied to explore aspects of cortical organisation where the application of functional methods is limited. Further applications may explore SAF topology itself in visual and auditory cortical areas of the congenitally blind and deaf as well as somatosensory cortex of individuals with tetraplegia, for which fMRI studies have shown preserved cortical retinotopic [48], tonotopic [49] and somatotopic [50] organisation, respectively.

The biological plausibility of fibre dispersion captured by sCF sizes demonstrates its applicability to explore detailed aspects of cortical hierarchical processing across the brain. It can be applied to study the cortical integration properties of SAF connecting, for instance, different levels within the auditory processing hierarchy [51], which have been shown to process different aspects of sound stimuli [45]. It can also be applied in the sensory-motor integration stream where population receptive field sizes in the primary somatosensory cortex (S1) have been found to be smaller than those in the primary motor cortex (M1) and, additionally, increase from the cortical representation of the thumb to that of the little finger [52]. Moreover, our approach is not restricted to a Gaussian model to accurately map the centre and size of the sCF, giving it broader applicability.

Although not directly comparable, our results are plausible given previous *post mortem* investigations that quantified connectivity in the early visual cortex of non-human primates. Tracer injections in macaque found feedback projections from V2 to V1 to cover 6.8 *±* 0.4 mm on the cortical surface [6]. The same study estimated the size of V1 to V2 feedforward projections in units of visual angle as 2.9 *±* 0.4^*°*^ [6]. These measures agree with our findings (Fig. 5a,d and Supplementary Fig. 12a,d) to within an order of magnitude, but cannot be directly and quantitatively compared to our metrics in the human brain. First, DWI tractography cannot distinguish feedforward and feedback connections and our metric is a weighted sum of both (see Appendix A). Second, tracer studies measure the contributions of axonal dispersion and telodendritic branching while DWI tractography captures only the axons. Third, direct transfer of findings from animal studies is not possible because of the significant expansion in human brain size and differences in gyrification patterns. Therefore, to directly relate the quantitative metrics, our method may be applied in animal brains in combination with tracer studies.

### Limitations

Our approach inherits the general limitations of DWI tractography and fMRI retinotopic mapping. Despite the improvements, biases of tractography remain important obstacles in mapping and interpretation of sCF estimates *in vivo*. We observed poorer performance of the method in the polar angle direction, likely explained by the gyral bias effects of tractography [22, 23] in the corresponding anatomical direction. Gyral bias becomes apparent in two ways. First, it causes a shift in the cortical terminations of SAF by generating a tendency to penetrate the gyral crowns (Fig. 6a,b) which, in our experiments occupied approximately 1/3 of the cortex in the polar angle direction. Second, the dominant long-range pathways terminating in V1 showed an almost reciprocal termination pattern in the cortex (Fig. 6c,d) likely preventing the detection of SAF expected to connect the walls of gyri/sulci (see Supplementary Fig. 13 for more detail).

Higher resolution DWI and advanced modelling may improve cortical sampling of the SAF by better disentangling the crossing fibre populations in the superficial white matter. Despite not resolving the methodological gyral bias, surface-based tractography approaches (e.g. [53]) may offer improvements especially in applications combining SAF connectivity with cortical maps on the surface.

Tractography-based sCF size estimates may depend on the modelled fibre orientation distribution and tractography algorithm. Also, as mentioned above, DWI-based metrics cannot distinguish feedforward and feedback connections, meaning that their contributions to the mapped sCF cannot be separated.

We used previously reported fMRI-estimated measures of cortical magnification factors and connective fields to interpret the mapped sCF sizes. Further experiments directly comparing structural and functional measures of connective fields sizes in the same individuals would help to more precisely characterise the relationship between structural and functional measures of cortico–cortical integration.

In conclusion, we have introduced and applied a novel method to characterise a prominent component of the human connectome: direct cortico–cortical connectivity by SAF. The method combines ultra-high spatial resolution DWI tractography with cortical topology mapped by fMRI to quantify the sCF, which reflect cortico–cortical information transfer and integration mediated specifically by the SAF. We show that SAF connecting different levels of the cortical visual hierarchy preserve the topological structure of cortical functional neuroanatomy. Our approach can be applied to investigate cortical organisation and reorganisation in topologically organised areas including the motor, somatosensory and auditory cortices. It may even be applied to explore and infer topological features in cortical areas whose topological mappings are unknown, as long as the limitations of tractography are borne in mind. The level of detail achieved by non-invasive mapping of SAF holds promise for applications previously only accessible by invasive tract tracing studies in animal research, opening new avenues for better understanding the human connectome.

## STAR Methods

The data used in this work were acquired as part of our previous study described in [21] in which details of data acquisition and preprocessing are also presented. Here, we briefly present the participant and data specifications and provide details of preprocessing and analysis used to support the sCF mapping framework.

### Participant information and data acquisition

Briefly, high spatial resolution DWI of a slab covering the occipital cortex were acquired (0.8 mm isotropic spatial resolution, b-values 800 and 1800 s/mm^2^, 60 diffusion-encoding directions per diffusion-weighted shell, 62 oblique near-axial slices originally placed to cover the V1 and V2 cortex, 25 interleaved non-diffusion-weighted volumes, anterior–posterior phase encoding) on a 3 T Connectom MRI scanner (maximum gradient amplitude 300 mT/m, Siemens Healthcare, Erlangen; applied maximum diffusion encoding gradient amplitude 188 mT/m) [54, 55] using a 32-channel radio-frequency head coil and a single-shot 2D spin-echo echo planar imaging sequence. Ten additional *b* = 0 s/mm^2^ volumes acquired with reversed phase encoding in the posterior-anterior direction were used for susceptibility-related distortion correction. Total acquisition time was approximately 45 min.

Functional MRI (fMRI) retinotopic maps were acquired at 1.0 mm isotropic resolution on a 7 T scanner (Terra Magnetom, Siemens Healthcare, Erlangen) based on a phase-encoded visual stimulation paradigm [26, 27]. The visual stimulation consisted of a contrast-reversing checkerboard stimulus restricted to expanding/contracting rings and a clockwise/anticlockwise rotating ray along the visual field eccentricity (Fig. 2a(i)) and polar angle (Fig. 2a(ii)) encoding directions, respectively [21]. T1-weighted structural MRI were acquired at 0.7 mm isotropic spatial resolution also at 7 T to enable mapping of V1, V2 and V3 on the cortical surfaces reconstructed with FreeSurfer [30].

Because the acquisition in [21] was optimised for V1 and V2 mapping, the acquired DWI slab covered V3 in only a subset of the data corresponding to 18 hemispheres (9 right, 9 left) from 11 participants.

## Functional MRI

### Smoothing the retinotopic maps

Functional retinotopic maps, polar angle (*φ*) and eccentricity (*ϱ*), were smoothed on the cortical surface to mitigate the effects of noise from fMRI acquisition on the fine-grained analysis required for sCF mapping. First, we determined for each vertex on the cortical surface if it corresponded to reliable functional signal. This was determined based on an arbitrarily defined signal-to-noise ratio (SNR) threshold of 5 computed as the magnitude of the fMRI signal at the stimulus frequency divided by the standard deviation of the frequency spectrum of the fMRI time-series. Second, for each reliable vertex we computed the mean retinotopy (*ϱ* and *φ*) value over a local neighbourhood [56] of reliable vertices defined around it. The identified unreliable vertices were omitted from computation of the mean. We repeated the process 4 times to achieve sufficient smoothing, determined by visualisation of the smoothed retinotopic maps.

### Delineating V1, V2 and V3 functional borders

Functional maps of V1, V2 and V3 were delineated manually on inflated cortical surfaces using FreeSurfer ‘TkSurfer’ based on the retinotopic maps for each hemisphere, separately. Cortical areal borders separating V1–V2 and V2–V3 correspond to the vertical and horizontal meridians of the visual field, respectively. V1, V2 and V3 were delineated by carefully drawing a line linking all points along the respective meridians by following the iso-eccentricity and iso-polar angle lines.

## Diffusion MRI

### Mapping fibre pathways using probabilistic tractography

Probabilistic streamline tractography [57] on fibre orientation distribution maps [58] was used to capture the dispersion of SAF pathways connecting V1, V2 and V3 using MRtrix3 [59]. We used cortical seeding to enable a high tracking seed resolution of 4 *×* 4 *×* 4 per voxel and at the same time, significantly reducing the resulting tractogram size compared to whole brain seeding. The seed region was created based on the V1, V2 and V3 cortical maps transformed to DWI volume space. Other tracking parameters were: curvature threshold of 30^*°*^, seed and tracking orientation distribution function (ODF) amplitude threshold of 0.1, tractography step size of 0.2 mm and reconstructed streamline lengths of 3–120 mm.

### Mapping SAF connections up to V3

SAF connecting V1, V2 and V3 were detected in surface space instead of the DWI volume space used in [21] to enable more precise mapping. The transformation of cortical maps from surface to DWI volume space for connectivity mapping creates overlapping regions between adjacent cortices. This makes the assignment of streamlines to each cortex ambiguous and thus, affects precise SAF mapping [21]. On the cortical surface, however, no overlap was introduced during manual delineation of the cortical maps. We transformed the streamlines generated by tractography from DWI to surface space using a previously computed non-linear deformation field between the DWI and structural MRI volumes using the ANTs software [60]. In surface space, the streamline coordinates were precisely mapped onto the cortex using an in-house python script [https://github.com/dchaimow/streamline_positions_from_surfaces/tree/v1.0.0]. For each streamline, the vertices surrounding its first and last cortical coordinates in the triangular surface mesh, taken as the streamline’s true cortical terminations, were identified. A streamline was considered to connect a pair of cortical areas *V*_*m*_ and *V*_*n*_ (*m< n*) if all the vertices surrounding its first and last cortical coordinates were a subset of the vertices of the respective cortical maps. To avoid bias from streamlines running entirely within the cortex, we removed streamlines with more than 80% intracortical length.

### Combining SAF with retinotopic maps

Combining retinotopic maps with SAF streamlines is an essential step in our sCF mapping framework. We used the distribution of the visual field coordinates, measured by fMRI retinotopic mapping, of SAF streamline terminations on the cortical surface to estimate the sCF. In surface space, each vertex is associated with a pair of visual field coordinates corresponding to smoothed eccentricity (*ϱ* = [1–6^*°*^]) and polar angle (*φ* = [*-*90^*°*^, +90^*°*^]). We assigned to each streamline the mean value of the visual field coordinates of the 3 vertices surrounding its first and last cortical coordinates, resulting in four visual field coordinates (*ϱ*_*m*_, *φ*_*m*_) and (*ϱ*_*n*_, *φ*_*n*_) per streamline connecting *V*_*m*_ and *V*_*n*_ (*m< n*).

### Assessing retinotopic order of SAF using tractography

To assess the detailed retinotopic organisation of SAF connecting *V*_*m*_–*V*_*n*_ (*m < n*), we created scatter plots of the visual field coordinates of their streamline terminations in *V*_*m*_ and *V*_*n*_ cortex, expressed in visual field units. Correlations for eccentricity (*φ*_*m*_, *φ*_*n*_) and polar angle (*ϱ*_*m*_, *ϱ*_*m*_) were calculated separately and illustrated as density plots over the entire group of 18 hemispheres, with the horizontal and vertical axes corresponding to visual field coordinates in *V*_*n*_ and *V*_*m*_, respectively. To enable combining the results across the left and right hemispheres, polar angle values in the left hemisphere were reflected across the horizontal meridian (see Fig.3 insert for polar angle) prior to generating the density plots. Pearson’s linear correlation coefficient (*ρ*) was computed between the visual field coordinates mapped in *V*_*m*_ and *V*_*n*_ by the SAF to quantify their expected retinotopic order mapped by tractography.

SAF retinotopic order also implies that retinotopy in a higher visual cortex (*V*_*n*_) can be predicted based on its SAF connectivity to a lower visual cortex (*V*_*m*_). To assess this, for each streamline connecting *V*_*m*_–*V*_*n*_ (*m < n*), the retinotopy values of the vertices representing its termination in *V*_*m*_ (*ϱ*_*m*_, *φ*_*m*_) were projected onto the vertices representing its termination in *V*_*n*_, thus, creating a SAF-generated retinotopy map in *V*_*n*_. The maps generated for each hemisphere were transformed to fsaverage surface space provided by FreeSurfer and averaged on a per vertex basis. They were compared against the corresponding group-averaged retinotopic maps obtained using fMRI. FreeSurfer atlas labels for V2 and V3 were used to restrict the visualisation to *V*_*n*_.

### Characterising sCF sizes mediated by SAF

The theory underlying sCF characterisation combining SAF tractography and fMRI retinotopic mapping is described in Appendix A. Experimentally, we estimated the sCF sizes by analysing the visual field coordinates of SAF cortical terminations for SAF connecting a defined neighbourhood in a higher visual cortex (*V*_*n*_) to the entirety of a lower visual cortex (*V*_*m*_). The neighbourhoods were defined by bins created according to the following procedure. In visual field coordinates, small bins of 1^*°*^ width were created in *V*_*n*_ in the cortical direction aligned with eccentricity encoding (Supplementary Fig. 11a,b). SAF terminating within each bin in *V*_*n*_ were identified and their visual field coordinates in *V*_*n*_, *x*_*n*_ = (*ϱ*_*n*_, *φ*_*n*_) as well as on their cortical terminations in *V*_*m*_, *x*_*m*_ = (*ϱ*_*m*_, *φ*_*m*_), were obtained. For each bin in *V*_*n*_, the distribution of the differences in the visual field coordinates, *x*_*m*_ *- x*_*n*_, was created separately for eccentricity (*ϱ*_*n*_ − *ϱ*_*m*_) and polar angle (*φ*_*n*_ − *φ*_*m*_) encoding (Supplementary Fig. 11c). The standard deviation of the *x*_*m*_ − *x*_*n*_ distribution for each bin was computed as a measure of the sCF size in the bin (Supplementary Fig. 11d).

Outliers, defined as data points beyond the [5 %, 95 %] percentile range of the distribution per bin, were removed for accurate estimation of the sCF sizes prior to computing the standard deviation. Additionally, polar angle values in the range [*−*90^*°*^, +90^*°*^] were converted from polar angle units to visual angle units (Fig. 2) by computing the arclength subtended by the polar angle at the corresponding eccentricity. The sCF sizes were reported in visual field (^*°*^) as well cortical surface (mm) units, the latter computed using the cortical magnification factors in *V*_*m*_ from [3], for eccentricity (*φ*) and polar angle (*ϱ*), separately.

## Acknowledgements

The research leading to these results has received funding from the European Research Council under the European Union’s Seventh Framework Programme (FP7/2007-2013) / ERC grant agreement n^*°*^616905. Funded by the Deutsche Forschungsgemeinschaft (DFG, German Research Foundation) – project no. 347592254 (WE 5046/4-2 and KI 1337/2-2). NW received support from the European Union’s Horizon 2020 research and innovation programme under the grant agreement No 681094, and from the Federal Ministry of Education and Research (BMBF) under support code 01ED2210.

## Author contributions

Conceptualization: F.M.A., E.K., N.W.; methodology: F.M.A., D.C., E.K.; software: F.M.A., D.C., D.H. for fMRI, K.P. for sequence development; formal analysis: F.M.A, E.K., D.C., L.J.E.; investigation: F.M.A., E.K., D.H.; resources: N.W., E.K., F.M.A; data curation: F.M.A, E.K.; Validation: F.M.A; writing original draft: F.M.A., E.K., D.C., L.J.E. and N.W.; review and editing: all co-authors; visualization: F.M.A and E.K.; supervision: E.K. and N.W.; project administration: F.M.A., E.K., N.W.; funding acquisition: E.K., N.W.

## A Appendix

### A.1 Theory of structural connective field mapping

We relate the novel structural connective fields (sCF) framework for characterising cortico– cortical integration by SAF to the established functionally defined connective fields framework that comprises an aggregate measure of cortical integration from intracortical and short-range white matter connections [3, 5]. First, we briefly describe the retinotopic processing of visual information in the early visual cortex and summarise the connective fields measures between different cortical hierarchy levels (V1–V2 and V1–V3) obtained in previous fMRI studies in the human brain. Second, we detail the theory underlying our *in vivo* method for sCF characterisation based on combining SAF streamlines from probabilistic tractography and the visual field coordinates of V1, V2 and V3 cortical areas from fMRI retinotopic mapping.

### A.2 Functional cortical connective fields estimated from fMRI

The retina samples the two-dimensional visual field parameterised by eccentricity (*ϱ*) and polar angle (*φ*) coordinates describing the distances (in degrees of visual angle) from the central fixation point subtended at the eye (Fig. 1a) such that two orthogonal directions are defined on the cortical surface.

The early visual cortex is retinotopically organised [18, 24–28]. This means that adjacent points in the visual field are encoded by neighbouring groups of neurons on the cortical surfaces of the hierarchically organised V1, V2 and V3. This organisational principle also imposes the two-dimensional visual field coordinate system – defined by eccentricity and polar angle – onto each cortical surface (Fig. 1a). Visual field coordinates can be mapped onto each location in V1, V2 and V3 using fMRI retinotopic mapping [26, 27]. We define *x*_*m*_ = (*ϱ*_*m*_, *φ*_*m*_) and *x*_*n*_ = (*ϱ*_*n*_, *φ*_*n*_) to represent a location on the visual cortex *V*_*m*_ and *V*_*n*_, respectively where (*m< n, m* = 1, 2, *n* = 2, 3).

The cortical representation of a unit (1^*°*^) of angular distance in the visual field is mapped by the cortical magnification factor (CMF) [3] into a distance along the cortical surface. CMF decreases as a function of eccentricity (*ϱ*), with more cortical space assigned to low eccentricities near the foveal representation than high eccentricities near the visual field periphery (Fig. 1a-ii and Supplementary Fig. 7b).

Although neurons representing a given location in the visual field respond maximally to stimuli at that location, they also respond to stimuli in the vicinity of that location, resulting in the integration of visual stimuli responses over a small region in the visual field. This region is known as the receptive field of the neuron.

It is not possible to measure the activity of individual neurons in humans non-invasively. To account for that, the receptive field concept has been extended to the population receptive field (pRF) [17]. The pRF describes the region in the visual field that stimulates an aggregate response from a local population of neurons, measured in an fMRI voxel. The pRF in a cortical area *V*_*n*_ is often modelled as a two-dimensional Gaussian function with standard deviation *o*_*V*_*n* defining the pRF size. The pRF size increases from the centre to the periphery of visual field and from low to higher hierarchy levels in the visual cortex from V1 to V3 (Supplementary Fig. 7a, the latter also in Fig. 1b).

Cortico–cortical integration between hierarchy levels in the early visual cortex has been estimated with fMRI using two different methods: mapping (i) cortical sampling size [3] and connective field size [5]. Both estimations are based on the analysis of aggregate functional activity, which reflects overall information integration comprising feedforward and feedback inter-area SAF as well as intracortical intra-area connections.

Approach (i) typically assumes a Gaussian model of pRF size such that pRF size in the higher area *V*_*n*_ can be computed by the convolution of pRF size in an earlier area *V*_*m*_ (*m< n*) and cortico–cortical sampling (cc-pRF) size. In this approach, cc-pRF size was determined in visual field coordinates as the square-root-difference of squared pRF sizes at different levels of the visual hierarchy and was converted to cortical coordinates in mm on the cortical surface using the CMF [3]. The cc-pRF sizes estimated using this method were approximately 3 mm for V1–V2 connections and increased along the visual hierarchy, reaching 5–6 mm for V1–V3 connections [3].

Approach (ii) typically defines the connective field of the higher area *V*_*n*_ as a two-dimensional Gaussian filter on the cortical surface of the earlier area *V*_*m*_ (*m< n*) that can optimally describe the temporal evolution of the cortical signal in *V*_*n*_. The width of the filter gives a measure of the connective field size. In agreement with the cc-pRF size estimates, connective field sizes obtained using this method were approximately 3 mm for V1–V2 connections and increased along the visual hierarchy, reaching 5–6 mm for V1–V3 connections [5].

### A.3 Structural connective field sizes estimated with SAF DWI tractography

Connectivity between two cortical visual areas *V*_*m*_ and *V*_*n*_ (*m < n, m* = 1, 2, *n* = 2, 3) can be characterised in terms of either divergence or convergence. Divergence for each finite neighbourhood around *x*_*m*_ in the lower area *V*_*m*_ is defined as the distribution of locations *x*_*n*_ in the higher area *V*_*n*_ connected to it. On the other hand, convergence for each finite neighbourhood around *x*_*n*_ in the higher area *V*_*n*_ is defined as the distribution of locations *x*_*m*_ in the lower area *V*_*m*_ connected to it. Either perspective, together with the total number of connections for each location, describes the connectivity between *V*_*m*_ and *V*_*n*_ completely. Here, directionality of connections is disregarded because DWI tractography cannot distinguish feedforward and feedback connections. Consistent with previous functional MRI studies, we use the convergence point of view in our analysis.

#### A.3.1 Definition of the structural connective field

Let us denote the number of axons connecting a finite neighbourhood around *x*_*m*_ in *V*_*m*_, to a finite neighbourhood around *x*_*n*_ in *V*_*n*_, as *N* (*x*_*m*_, *x*_*n*_). For simplicity, we assume that (i) the sizes (in retinotopic coordinates) of the neighbourhoods are equal, (ii) a finite set of points can be defined in each area such that their neighbourhoods fully cover the area without overlapping, (iii) the retinotopically matching set of these points are used in both areas, and (iv) there are a non-zero number of connections to *V*_*m*_ from each neighbourhood of *x*_*n*_ in *V*_*n*_.

For a particular neighborhood around *x*_*n*_ we can then define a normalised distribution describing the proportion of the number of fibres connecting the neighbourhoods around *x*_*n*_ and *x*_*m*_ relative to the total number of connections between the neighbourhood of *x*_*n*_ and the entirety of *V*_*m*_ (*N* (*V*_*m*_, *x*_*n*_)):

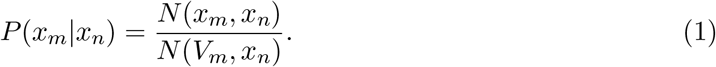

We assert that there exists an integrable probability density such that

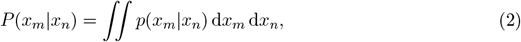

where the integrals are over the neighbourhoods of each point. This distribution describes the structural connective field of *x*_*n*_ to *V*_*m*_. In line with the retinotopic connectivity principle, we expect that the expected values of the structural connective field 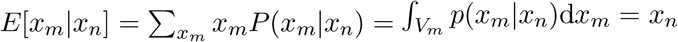, meaning the centre of the structural connective field is the retinotopically corresponding point on the cortical surface of *V*_*m*_. The standard deviation of this distribution *σ*_*mn*_(*x*_*n*_), defined in an analogous way to the expected values, is the measure of the structural connective field size (Fig. 2).

#### A.3.2 Contributions of feedforward and feedback projections

To relate the novel sCF concept to connectivity measures derived from tracer studies – where feedforward and feedback connections can be separately traced using retrograde and anterograde tracers – we derive the contributions of both feedforward and feedback connections to the sCF size below.

The total number of connections *N* (*x*_*m*_, *x*_*n*_) can always be decomposed into the sum of the number of feedforward *N*_*m→ n*_(*x*_*m*_, *x*_*n*_) and the number of feedback *N*_*n → m*_(*x*_*m*_, *x*_*n*_) axons:

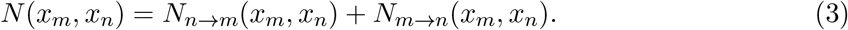

Feedback and feedforward fibres can be characterised by their corresponding structural connective fields 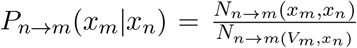 and 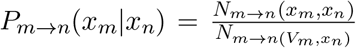 with sCF sizes *σ*_*n → m*_(*x*_*n*_) and *σ*_*m → n*_(*x*_*n*_), respectively. Then, the total structural connective field can be represented as a sum of the structural connective fields of the feedback and feedforward projections, weighted by their relative numbers:

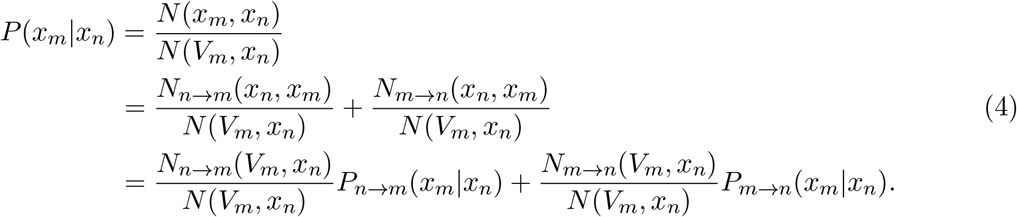

It can be shown that sCF, defined as *σ*_*mn*_(*x*_*n*_) of the distribution *P* (*x*_*m*_|*x*_*n*_), can be obtained from the sCF sizes of feedback and feedforward fibres *σ*_*n → m*_(*x*_*n*_) and *σ*_*m → n*_(*x*_*n*_) using the following relationship if the covariance is zero and the means of the two distributions are equal:

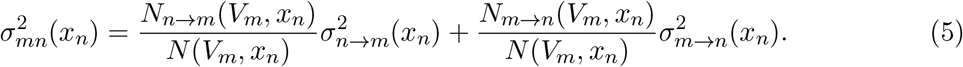

The assumption that the mean of the distributions for feedback and feedforward fibres are equal is justified by the retinotopic connectivity principle.

#### A.3.3. Estimating sCF sizes from SAF tractography

Assuming that the spatial distribution of streamlines mapped by probabilistic DWI tractography is unbiased and accurately reflects the underlying distribution of SAF connections, we can estimate the sCF size using the following strategy. For each streamline *i* connecting *V*_*m*_–*V*_*n*_ we can determine the visual field coordinates (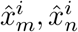) (each composed of polar angle *φ* and eccentricity *ϱ*) of its cortical terminations by sampling the cortical retinotopic maps obtained using fMRI at the streamline’s terminations on the *V*_*m*_ and *V*_*n*_ cortical surfaces, respectively (Fig. 2). These coordinates are given by:

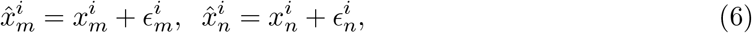

where (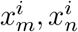) denote the true coordinates of axonal terminations, and 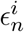 and 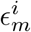 are the aggregate measurement error resulting from inaccuracies of fMRI retinotopy and DWI tractography, assuming that the noise is additive. We assume that tractography and retinotopy estimates are not biased (their errors have zero mean e.g. *E* [*ϵ*_*n*_]= 0), and that tractography errors and biological dispersion of the fibres in white matter are statistically independent.

Then, for each point 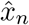 in *V*_*n*_ a structural connective field is estimated as a normalised distribution of the experimentally mapped visual field coordinates 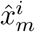 in *V*_*m*_ by the SAF streamlines. The estimated connective field 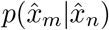 can be expressed as a convolution of the true biological connective field *p*(*x*_*m*_|*x*_*n*_) with the distribution of errors, which we assume to be Gaussian distributed and independent of *x*_*m*_ and *x*_*n*_ for simplicity:

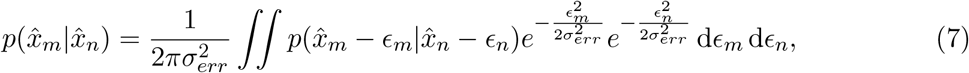

where 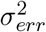 is the error variance. The estimated sCF size is a standard deviation of the 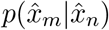 with respect to 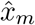. Since the second centred moment of the convolution of two functions is a sum of the second moments of these functions if at least one of them is a centred distribution, we can write

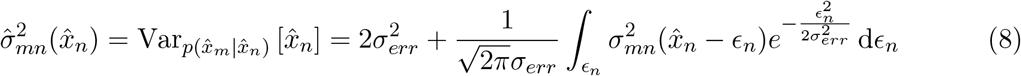

Thus, the estimated sCF size is a sum of the double error variance 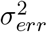 and the true biological sCF size, smoothed along the *x*_*n*_ coordinate by a Gaussian filter. If the error is small compared to the biological sCF size, Equation 8 provides an accurate estimate of sCF.

Since connective field sizes depend on eccentricity, we estimate the sCF sizes in clusters of SAF streamlines binned along the eccentricity encoding direction in *x*_*n*_ in bins of 1^*°*^ size. The binning procedure results in an additional boxcar average of the smoothed eccentricity-dependent measure of sCF size described in Equation 8.

