## Supplemental information for "Connectivity at fine scale: mapping structural connective fields by tractography of short association fibres *in vivo*"

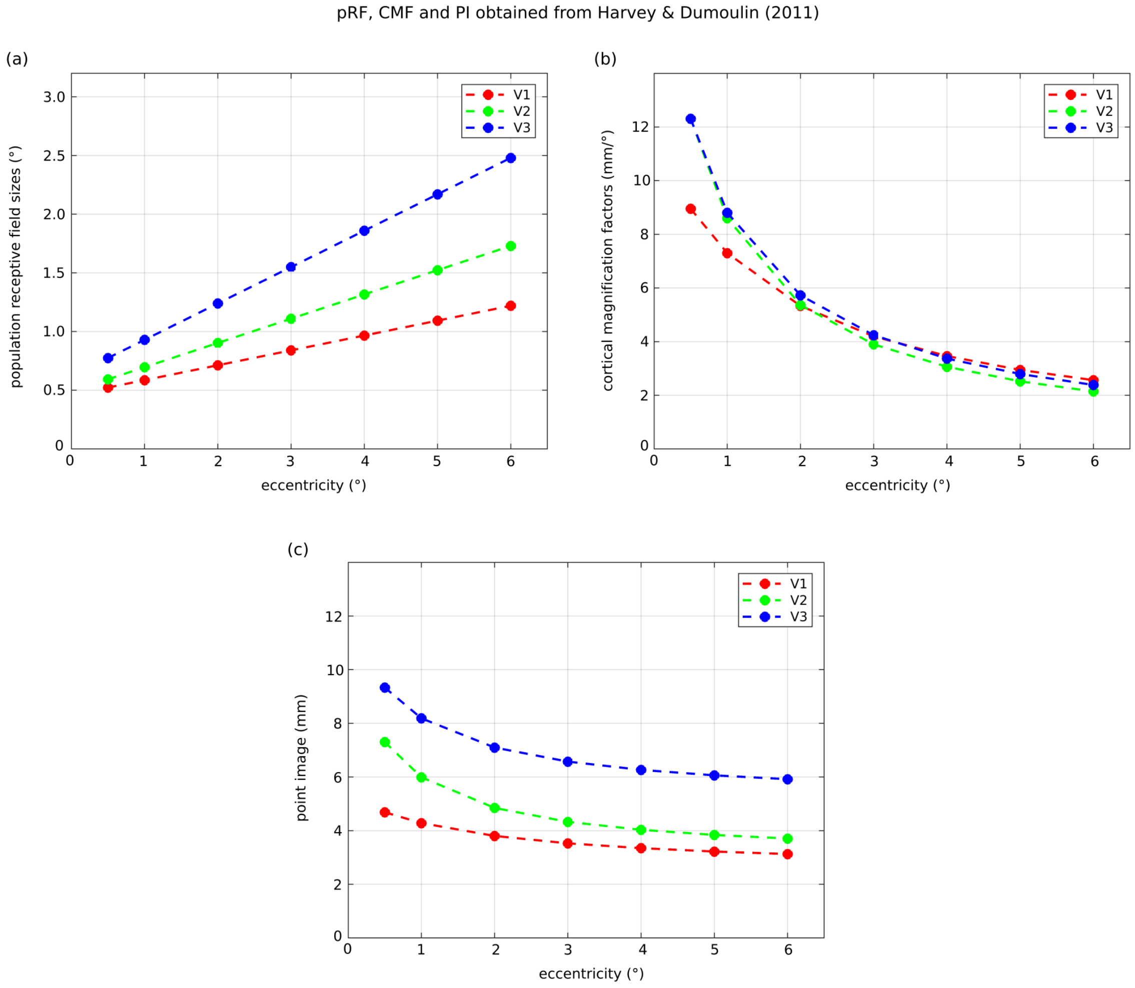


Fig. 7: Functional MRI estimates of (a) population receptive field (pRF) size, (b) cortical magnification factor (CMF) and point image size (pRF×CMF) as a function of eccentricity for V1, V2 and V3 [3].

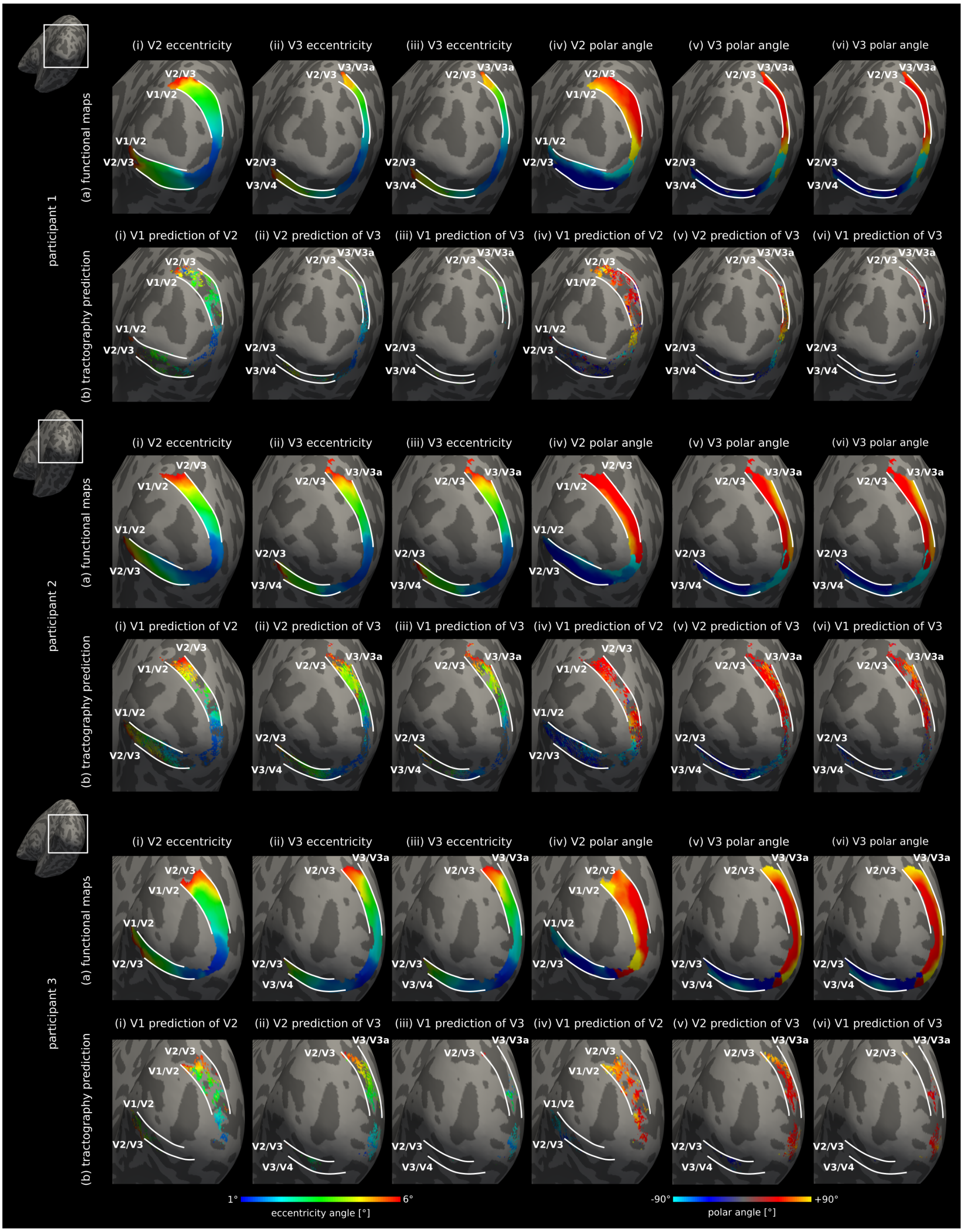


Fig. 8: Single-hemisphere predictions of cortical retinotopy based on SAF connectivity for participants 1–3. (a) Visual field coordinates – eccentricity and polar angle – in V2 and V3 determined by fMRI. (b) Corresponding tractography-generated predictions based on SAF connecting (i,iv) V1–V2 and V1 retinotopy, (ii,v) V2–V3 and V2 retinotopy, and (iii,vi) V1–V3 and V1 retinotopy. Each individual's cortical labels, delineated manually in native space, were used to determine the V1, V2 and V3 areal borders on the brain surface.

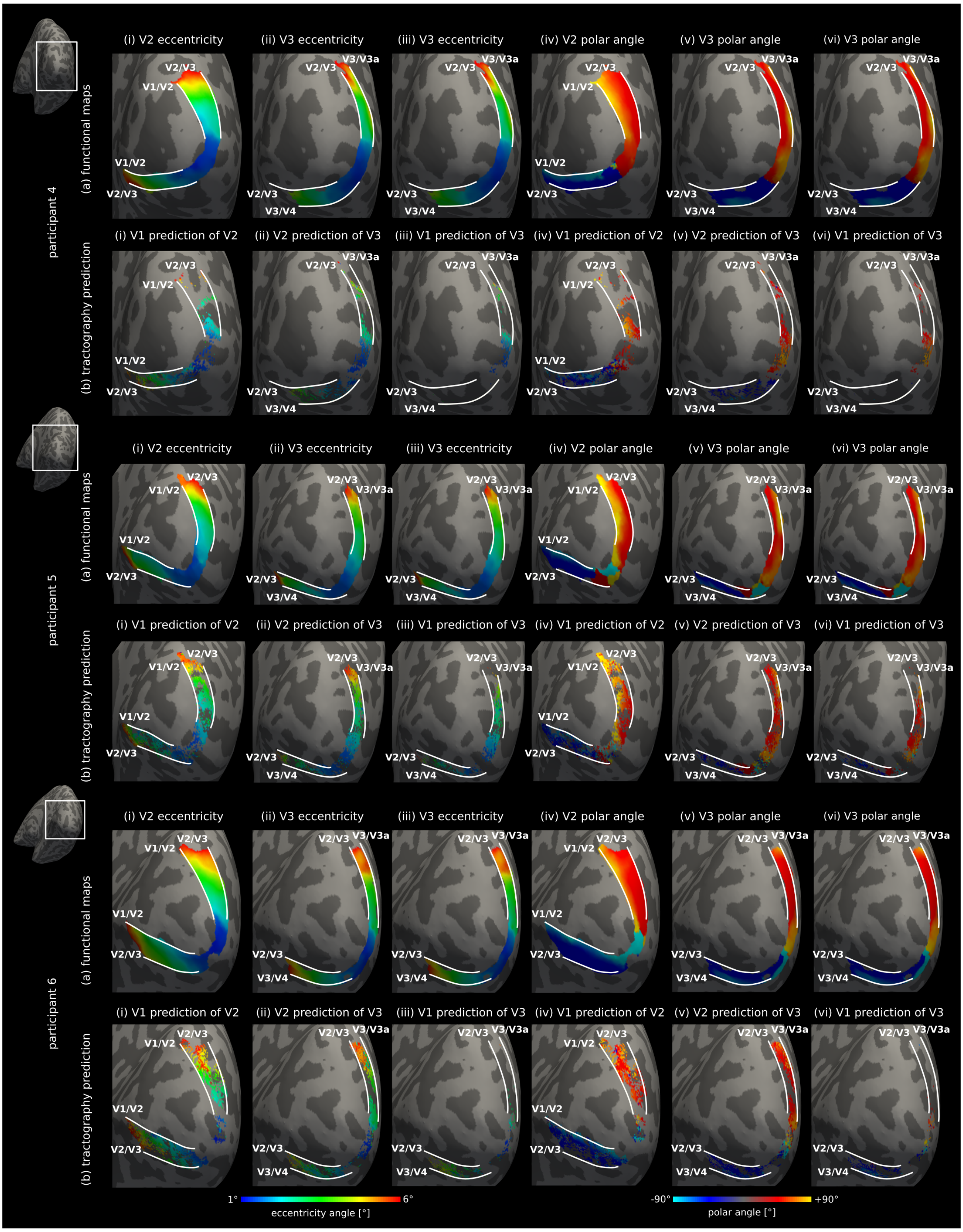


Fig. 9: Single-hemisphere predictions of cortical retinotopy based on SAF connectivity for participants 4–6. (a) Visual field coordinates – eccentricity and polar angle – in V2 and V3 determined by fMRI. (b) Corresponding tractography-generated predictions based on SAF connecting (i,iv) V1–V2 and V1 retinotopy, (ii,v) V2–V3 and V2 retinotopy, and (iii,vi) V1–V3 and V1 retinotopy. Each individual's cortical labels, delineated manually in native space, were used to determine the V1, V2 and V3 areal borders on the brain surface.

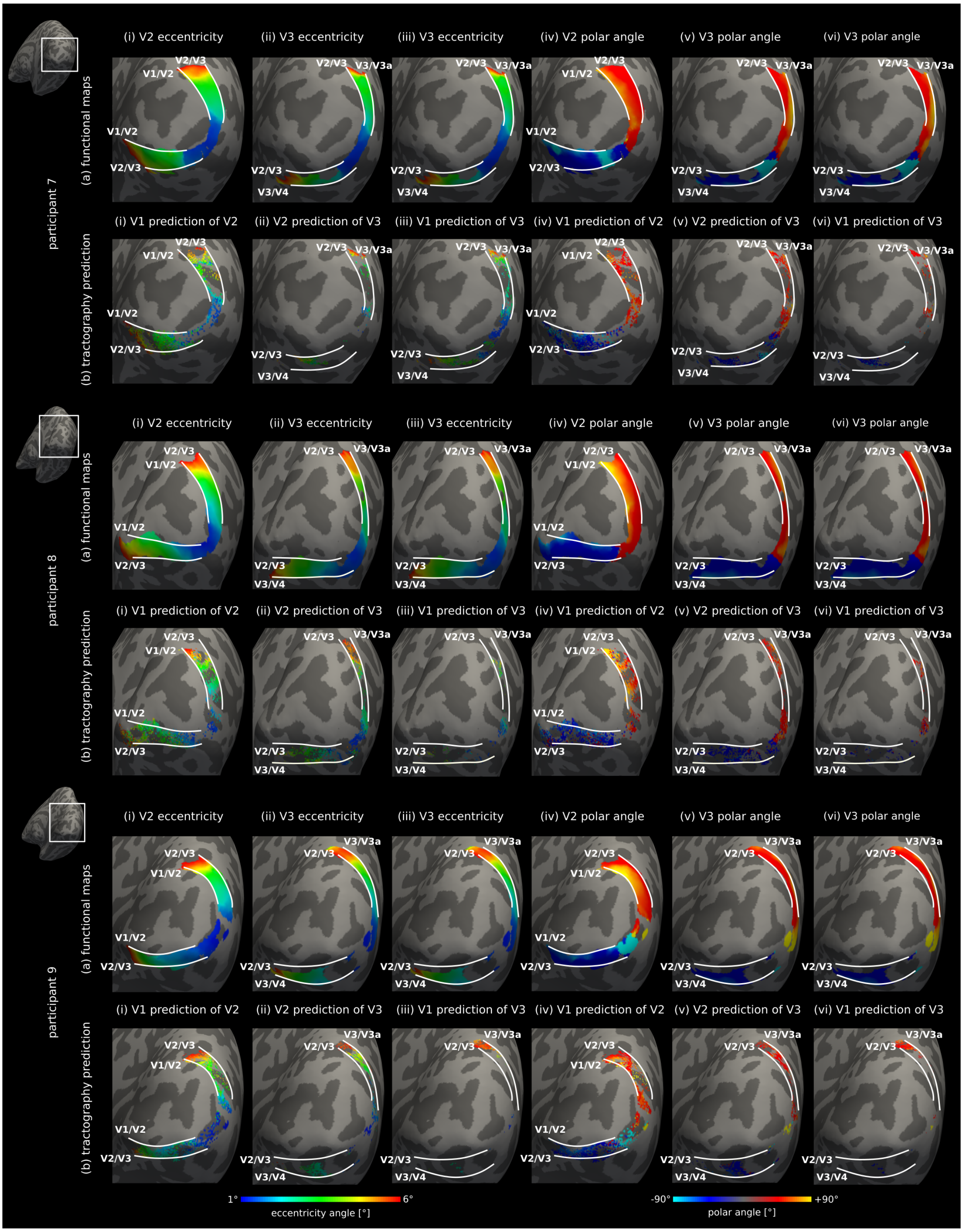


Fig. 10: Single-hemisphere predictions of cortical retinotopy based on SAF connectivity for participants 7–9. (a) Visual field coordinates – eccentricity and polar angle – in V2 and V3 determined by fMRI. (b) Corresponding tractography-generated predictions based on SAF connecting (i,iv) V1–V2 and V1 retinotopy, (ii,v) V2–V3 and V2 retinotopy, and (iii,vi) V1–V3 and V1 retinotopy. Each individual's cortical labels, delineated manually in native space, were used to determine the V1, V2 and V3 areal borders on the brain surface.

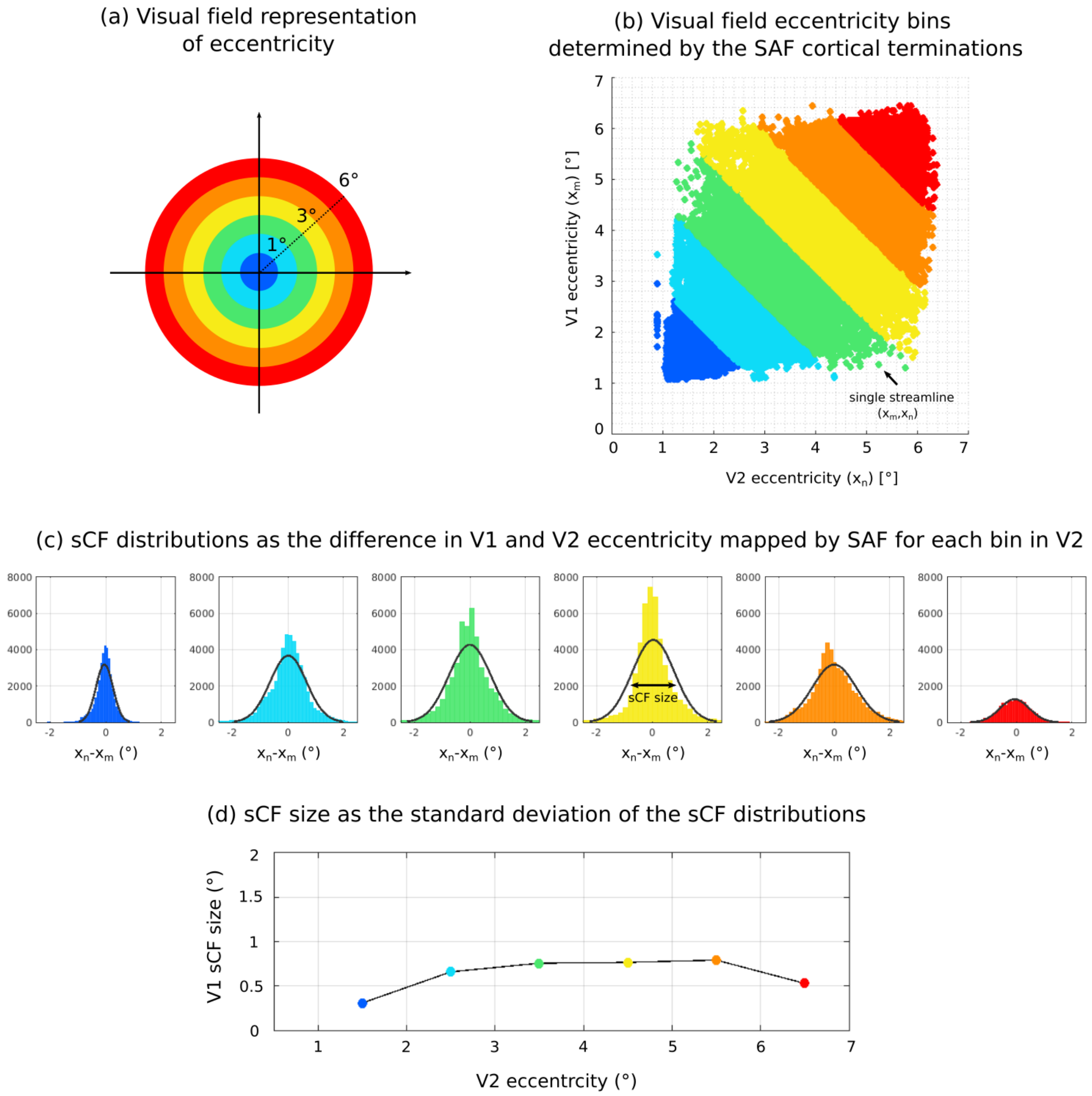


Fig. 11: Characterisation of structural connective fields (sCF) sizes mediated by SAF, illustrated here for V1–V2 SAF in the eccentricity encoding direction. (a) Visual field representation of eccentricity in the experimental range of approximately 1–6°. (b) Bins in eccentricity with a width of 1° were created in the higher cortical area V2 (or another higher area $V_n$). For each bin, cortical terminations of SAF streamlines in V1 (or another lower area $V_m$) were identified, each represented by one point on the scatter plot. Here, $x_m$ and $x_n$ represent the visual field coordinates of SAF streamline terminations on $V_m$ and $V_n$ cortical surfaces, respectively. The logarithmic scale reflects the cortical magnification factor in the respective cortices. (c) SCF was defined for each bin separately as the distribution of the differences in the sampled visual field coordinates ($x_2 - x_1$ or for other areas $x_n - x_m$) along eccentricity. (d) The sCF size was estimated as the standard deviation of the distribution in each bin. In (d) the centre of each bin is shown on the horizontal axis.

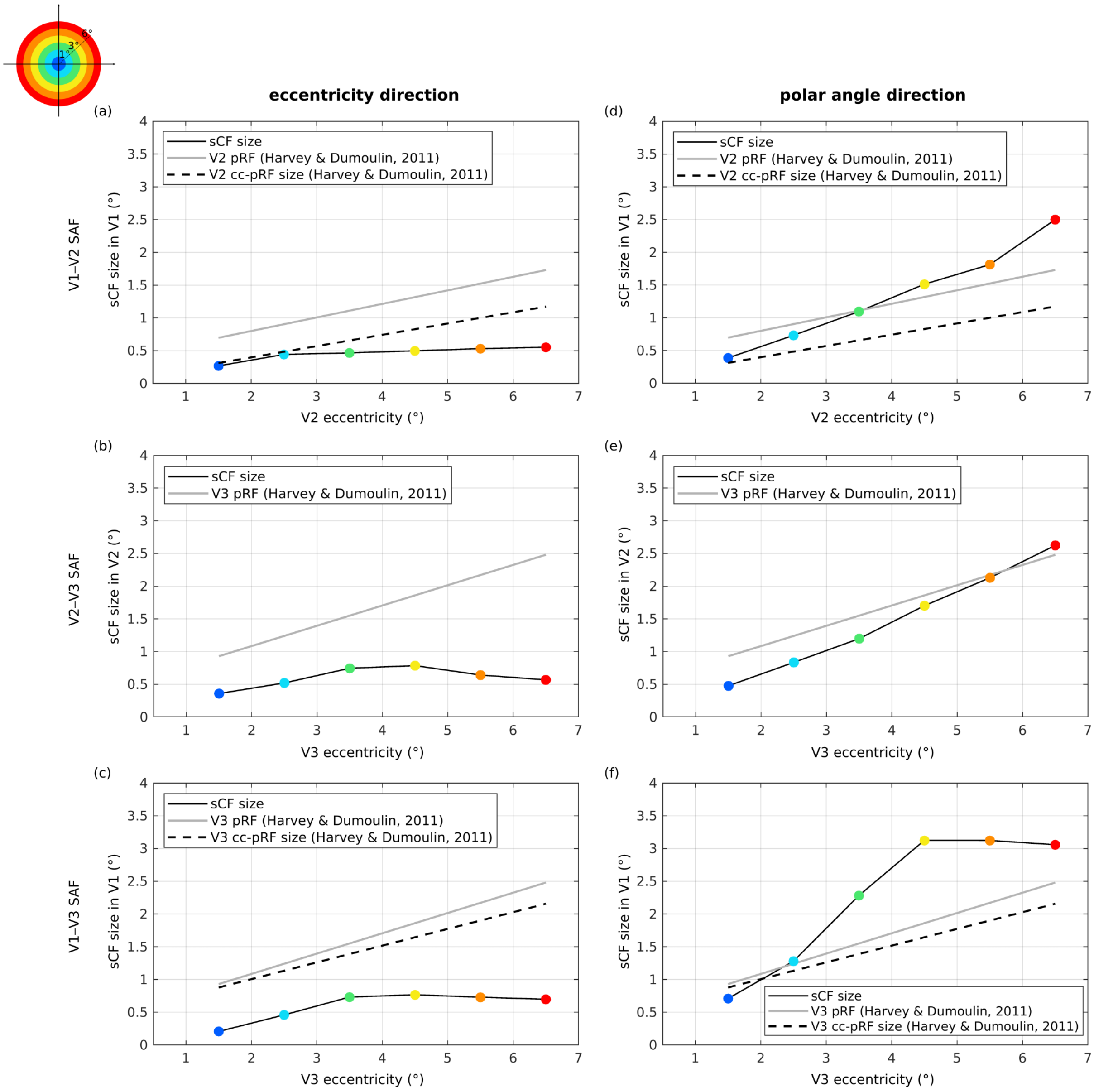


Fig. 12: Estimated sCF size (°) for cortical directions aligned with eccentricity (a-c) and (d-f) polar angle encoding. Group-level estimates are shown for V1–V2, V2–V3 and V1–V3 connections as black solid lines connecting the coloured points. The sCF size was estimated as the standard deviation of the distribution of the differences in the visual field coordinates of SAF terminations sampled on the cortical surface in $V_m$ and $V_n$. For both eccentricity and polar angle, the sCF size was calculated for SAF terminating in 1° bins of visual field eccentricity in $V_n$ ($m < n$). Each coloured point corresponds to the sCF size of one bin. For comparison, literature values (from Ref. [3]) of fMRI-estimated $V_n$ receptive field (pRF) sizes are plotted as gray solid lines and cortico-cortical sampling (cc-pRF) sizes for sampling of $V_m$ in $V_n$ are plotted as dashed lines (see Appendix A for definition of cc-pRF size). That work did not report cc-pRF sizes for V2-V3 connections, hence not shown.

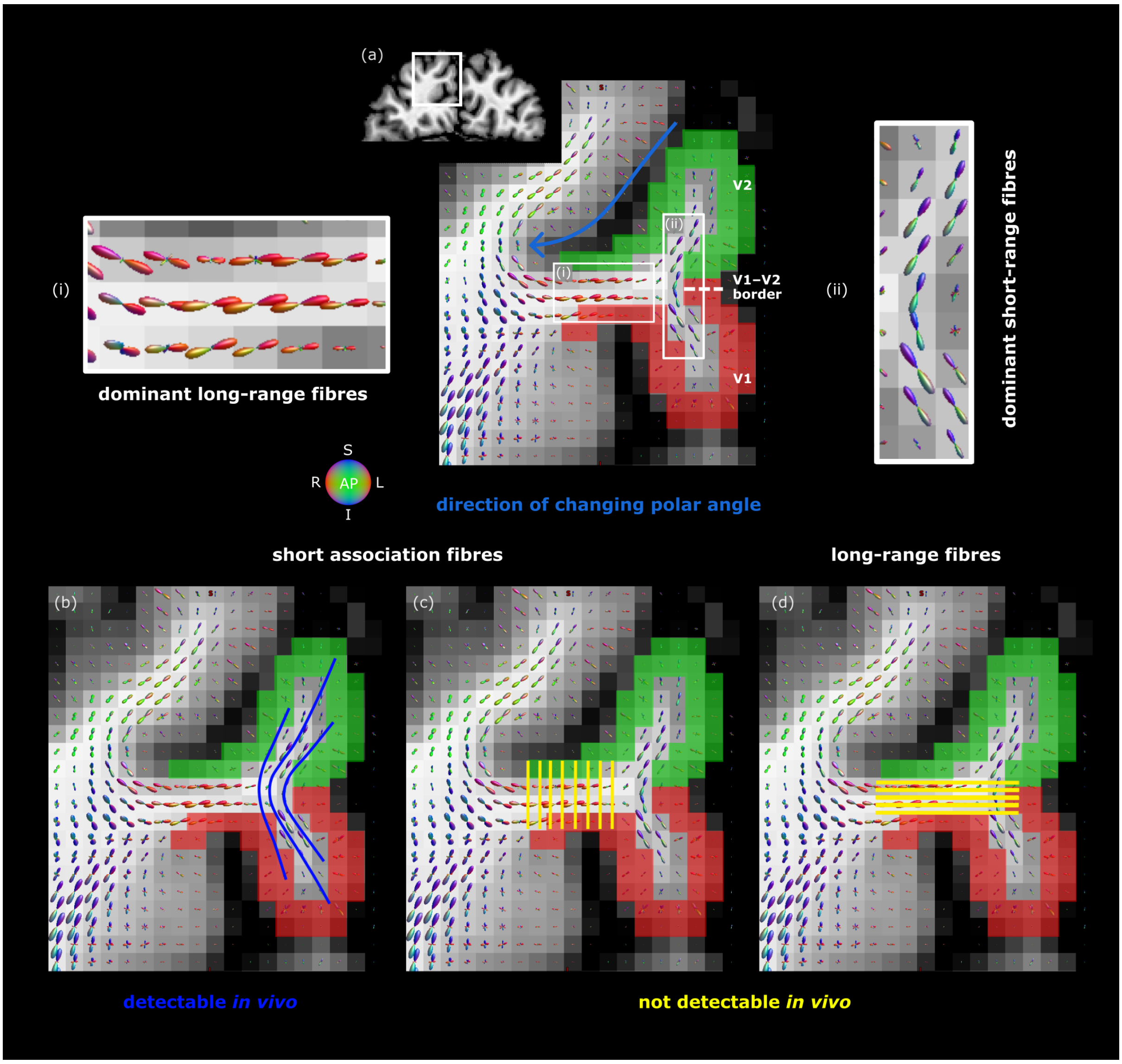


Fig. 13: Gyral bias of tractography in the polar angle encoding direction. (a) The interface of a V1 and V2 cortex is shown on a coronal plane in a representative participant. On the cortical surface, polar angle mapping corresponds to the anatomical direction with the largest variations in sulcal depth (blue arrow). Tractography is differentially affected in this direction because of gyral biases [22, 23]. (b,c) Not all SAF expected to connect V1–V2 could be detected by tractography. The detected SAF terminated mainly on and near gyral crowns (b), biasing their cortical coverage to regions best accessible by tractography and thus, resulting in the gyral bias. However, retinotopy expects V1–V2 SAF to also connect sulcal walls over short and almost straight pathways (c). These connections could not be mapped likely because of their crossings with the dominant long-range fibres (a(i)). (d) Similarly, the long-range fibres expected to penetrate the cortex at sulcal troughs could not be detected. This was likely because of their crossings with the dominant SAF in superficial white matter (a(ii)). Highly coherent and dense fibres demonstrated in the fibre orientation distribution maps in the locations of the (a(i)) long-range and (b(ii)) SAF pathways show their dominant arrangements, which appear to contribute to the observed gyral bias effect. This "winner-takes-all" principle possibly explains the observed gyral bias in the polar angle direction. A: anterior, P: posterior, I: inferior, S: superior, L: left, R: right.
